# Frequency-specific theta states in the hippocampus are linked to reconfiguration of population activity with respect to behavioural context

**DOI:** 10.1101/2024.12.11.627908

**Authors:** Laura Masaracchia, Pablo Oyarzo, Felipe Fredes, Diego Vidaurre

## Abstract

Neural activity reflects both external stimuli and the brain’s internal state, which shapes how information is processed and perceived. An example of modulation of neural responses by network states is phase-precession in the hippocampus, where the phase of theta oscillations affects the firing of single neurons (place cells) and its relation to the external world. Here, we examine a different form of relationship between oscillations and neural activity, where frequency and power of the oscillation, instead of phase, are associated with differential neural firing patterns at the population level. We refer to this as *ensemble pattern reconfiguration*. To study this effect, we use electrophysiological recordings of rats performing an odour-memory (non-spatial) task. Using a data-driven model, we identified two distinct theta states: low-power-lower-theta (LPLT) and high-power-higher-theta (HPHT). Through regression and decoding analyses, we found that these states are associated to modulations in hippocampal neural ensemble activity representing trial outcome - though the effect did not consistently reach statistical significance in all our subjects. Our findings suggest that power and frequency variations within theta oscillations may be linked to the reconfiguration of neural firing at the network level, motivating further investigation in larger datasets.

## Introduction

Even without being engaged in a task, an animal’s brain activity is in perpetual motion to shape its cognitive and physiological state ^1,2^. Any sensory stimulus coming from the external world is then met with an ever-changing landscape of neural activity, which inevitably influences the way a stimulus is processed and perceived ^3–5^. This means that even identical stimuli are processed differently, depending on the current brain state ^6–13^; and that brain states affect information processing at the neural level ^14–20^.

At the regional (mesoscopic) level, network states can be identified with oscillatory patterns in the local field potential (LFP) signal, often associated with cognitive and physiological functions ^21–27^. Although the LFP signal is typically attributed to synaptic activity, neural firing can affect these oscillations ^28^. In turn, these extracellular membrane fluctuations affect firing activity of the local neurons, through ephaptic effects ^29^, influencing cognition, and suggesting that the LFP oscillatory patterns may reflect a functional purpose ^30,31^. Existing literature targeting the relation between oscillations and neural firing has traditionally focused on the modulation of neural firing rate by the phase and amplitude of oscillations, across animals, brain regions, frequency bands and tasks ^32–38^. Although amplitude and frequency of oscillations are known to be related ^39^, less focus has been devoted to the link between frequency of oscillations and neuronal firing ^40–42^. Existing work in this regard investigated fluctuations in peak frequency of oscillations and their effect in modulating neural excitability^43^.

An example of oscillatory patterns associated to cognitive functions is theta oscillations in the hippocampus, studied within memory formation and recall, and spatial navigation ^21,26,27,44–47^. A well-established example of oscillation-to-neuron modulation is given by the phase precession phenomenon, where hippocampal place cells fire at specific phases of theta oscillations, depending on an animal’s location in space ^48^.

In this study we explore a kind of relationship between oscillations and neuron activity in the hippocampus, where variations in frequency and amplitude of theta oscillations (here associated with distinct states) are linked to differential representations of behavioural information (here whether the trial was performed successfully or not) at the neural population level, in rats performing an odour-memory, non-spatial task. To do so, we use a combination of data-driven methods for state detection and decoding analysis to link the detected states to neural activity patterns. First, we find two distinct states within the classically defined theta band: low-power-lower-theta (LPLT) and high-power-higher-theta (HPHT). Then, we quantify the difference in the neural ensemble patterns reflective of trial outcome between the two theta states, through prediction and decoding analyses. Our results suggest a tendency for neural population activity to differentially represent success and failure depending on which theta state is active. This suggests that the hippocampus may use different network configurations to perform different functions with the same neural population. Of note, our analyses do not necessarily establish a causal link from the LFP to the spikes (or the other way around), but an effect across scales (meso-scale in the LFP signal and microscale in the neural spikes).

## Results

In this study, we quantified the relation between mesoscale network states (represented as oscillatory patterns) and how neural firing patterns relate to behaviour. For this, we used previously recorded local field potentials (LFP) and spike data from the dorsal CA1 region of the hippocampus of four rats performing an odour-memory, non-spatial task ^49,50^. The task required rats to recognise whether presented odours followed the correct order within a previously learned sequence. Correct responses were rewarded with water after each odour presentation. We have introduced **Figure 1** to schematically visualize our method pipeline, that we summarize below. This is meant as a reference to aid the understanding of the method, rather than as a figure reporting results. We investigated whether the CA1 neural population reflected differently trial outcome depending on the local network state (**Figure 1A**). For simplicity, we limited our analysis to two states, without placing a strong interpretation on this number (also, see **Methods** about practical considerations in the choice of the number of states). We fitted hidden Markov models (HMM ^51–53^, a data-driven method) to detect the hippocampal states from the LFP data. The HMM assigned a state to each time point. We then linked these states activations to the corresponding time points in the spike data (**Figure 1B**). With this information, we performed two different analyses: neuron-state interaction analysis, quantifying the effect of the interaction between the oscillatory states and neural activity in predicting trial outcome (**Figure 1C**); and state-conditioned decoding analysis, quantifying the difference in neural population patterns representing success vs failure between states (**Figure 1D**).

**Figure 1:**
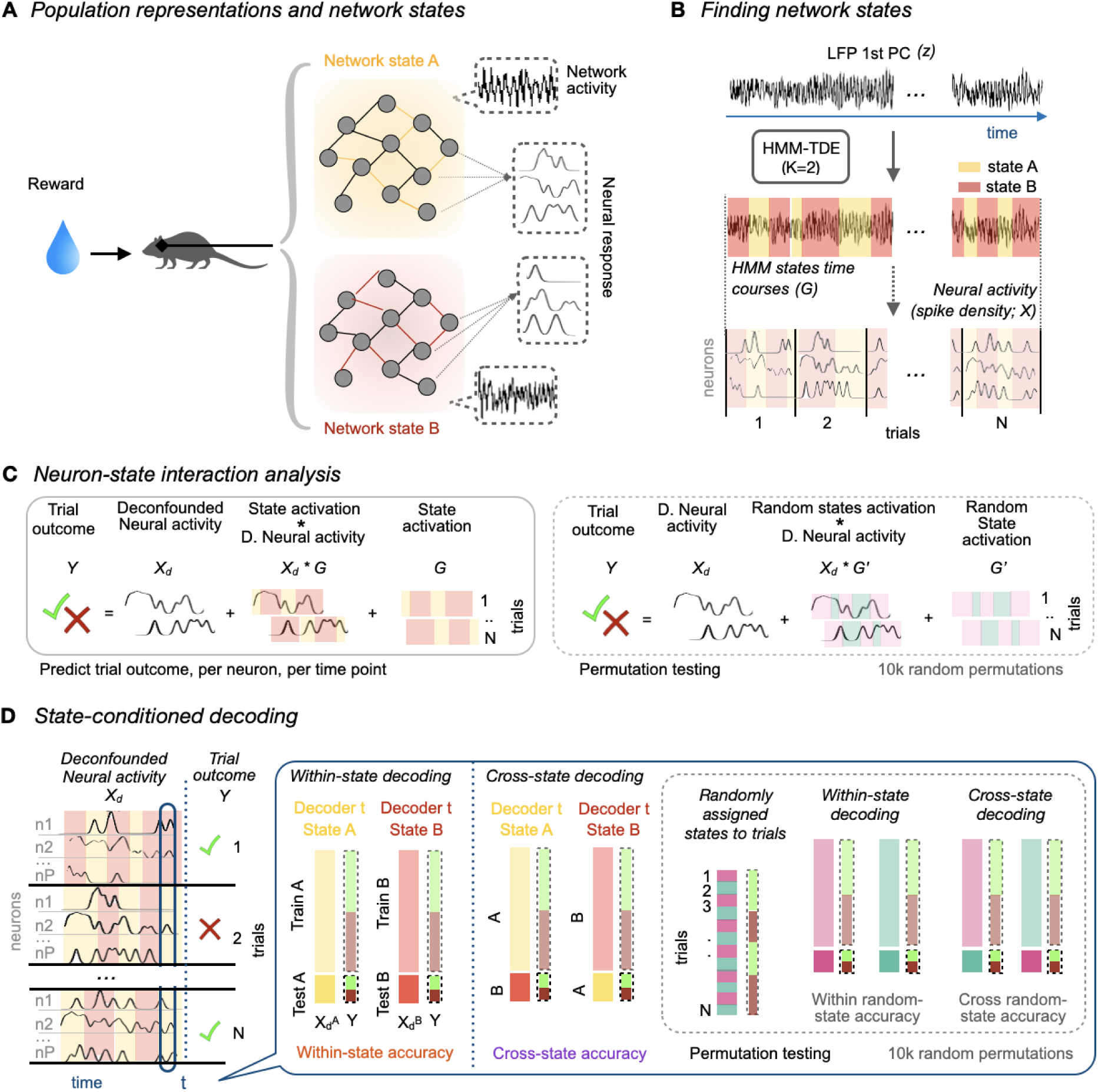
Schematic illustration of the hypothesis and methods. **A.** *Population representations and network states.* Neural population representation of an input depends on the state of the network in which the neurons are embedded. Here we show a visual example of our underlying hypothesis: dorsal CA1 neurons’ response to an input (for example, reward after a successful trial) varies with the network configuration (e.g., state A or B). Network activity is here represented by the LFP signal. **B.** *Finding network states:* an HMM with K=2 states is run on the continuous LFP data (*z*; top row). The HMM assigns a state to each point of the LFP data (state time courses, *G*, with the signal in middle row). The state time courses are overlied on the neural activity (spike density, *X*), associating the activity of the neural population at each time point to a state. All the data are then epoched into trials. **C.** *Schematic of the neuron-state interaction analysis*: we predict trial outcome (*Y*) from three predictors: neural activity deconfounded for motion (*X_d_*), state activation (*G*) and the neuron-state interaction (the product of deconfounded neural activity and state activation, *Xd*G*), per neuron and per time point. The states contribution is quantified by means of permutation testing, comparing the prediction error of this model with another using random states (G’; 10,000 random permutations). **D.** *Schematic of state-conditioned decoding:* for each time point, trials of deconfounded neural activity (*Xd*) are grouped into two sets, depending on the state activation at that time point (e.g., state A or B). Then, a decoder is used to predict trial outcome (*Y*) from each state-related set of trials (and for each time point), independently, in a custom train-test, class-balanced decoding procedure. We perform two measures, a within-state prediction (from a decoder trained and tested on the same state), and a cross-state prediction (from a decoder trained on one state and tested on the other). These two measures are then compared to the analogous analysis performed on randomly assigned states, with 10,000 random permutations.

### Low-power-lower-theta (LPLT) and high-power-higher-theta (HPHT) hippocampal states

Instead of prespecifying the definition of network states in terms, for example, of frequency bands, here we used a data-driven, unsupervised model (the HMM ^53^) to detect states according to the LFP’s spectral features ^54^. The states time courses output by the HMM are a time-point-by-time-point binary classification of the continuous data into two different groups (states), where each group reflects differences in frequency and power within the data. The HMM, run on the continuous broadband LFP data (**Figure 2A**), identified one state with low power and a major peak in the lower theta (7-8 Hz) frequency band (LPLT), and another with high power (1.2 to 3 times higher than that of LPLT) and a major peak in the higher theta (8-9 Hz; HPHT). As the HMM was run on the broadband signal, the states’ power spectra comprised multiple frequency bands. We visualized the ratio between the two states’ power spectra (**Supplementary Figure 1**) to check for proportional differences in the power spectrum across the broadband frequency range and confirmed that the largest difference between the two states was within the theta frequency range, consistently across all subjects. **Figure 2B** shows the states’ power spectra. We then compared statistically power and frequency across states, finding significant differences in both power and frequency distributions in three out of four subjects (**Supplementary Figure 2).**

**Figure 2:**
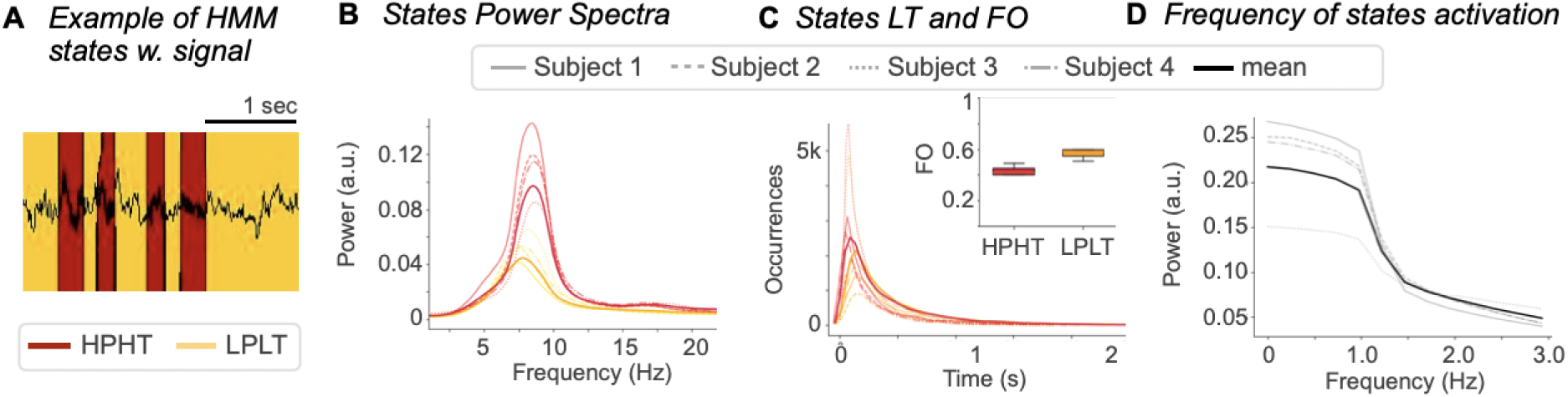
Characterisation of the HMM states. **A.** *Example of HMM states with signal*. An example of the HMM state assignment of the LFP data, for one subject. **B.** *States Power Spectra.* Frequency and power content of the two states for each subject (transparent lines, different linestyles per subject). The HMM consistently finds a high-power, higher theta (HPHT, red) and low-power, lower theta (LPLT, yellow) state across subjects, even if run independently for each subject. Here reported also each state mean across the four subjects (thick darker lines). **C**. *States lifetime (LT) and fractional occupancy (FO)*. The lines show the length in seconds of each state occurrence, per state and per subject (transparent lines). HPHT and LPLT have similar lifetime distributions, with LPLT states typically being active slightly longer (yellow distribution is lower and peaking more right than red distribution). Also shown the mean lifetime across all subjects (thick darker lines). The box plot shows fractional occupancy (FO), the proportion of time of activation of each state, across subjects. LPLT is typically active around 60% of the time, HPHT, around 40%. **D.** *Frequency of states activation*. Shown for each subject how often there is a state switch (gray lines). Most state switches occur with a frequency below 2Hz, meaning at most two switches per second. Also shown the average across four subjects (black thick line).

As a sanity check, we tested the sensitivity of the HMM to small spectral changes. To do so, we run the HMM on simulated brain data, varying only in frequency between 7 to 9 Hz. The model successfully distinguished two states based on these small frequency differences (**Supplementary Figure 3**).

**Figure 2C** shows the states lifetime (LT, the average duration the state activations) and fractional occupancy (FO, defined as the proportion of time that a state was active in total). Both temporal (e.g., most states activations lasting between 0.1 and 0.8 seconds) and spectral (e.g. the theta dominance) features suggest that neither of the two states represents a motion artefact. To investigate whether the states were linked to different phases of theta, we computed the frequency of their activation. If the states switched with a frequency between 7 and 9 Hz, they could have been linked to the phase of the oscillation. We found that the states reoccurred predominantly with a frequency of 0.1 to 1.2 Hz, indicating no direct link to the theta phase (**Figure 2D**). To explicitly test the link between states and theta phase, we statistically compared the distributions of theta phases between the two states (see **Methods** for more details). Our analysis revealed that the two states did not have a significantly different distribution over theta phase in two out of four subjects (Subjects 1 and 4, where the difference in mean value of the states’ phase distributions was below 0.01 π, and the difference in variance below 0.005 π; the p-value of the Watson U^2^ test, corrected for multiple comparison with Benjamini-Hochberg method, was not significant), and small but significantly different theta phase distributions in the other two subjects (Subjects 2 and 3, with a difference in mean value of the distributions of 0.04 π and 0.06 π respectively, and difference in variance of 0.03 π; p-value of the Watson U^2^ test, corrected for multiple comparisons with Benjamini-Hochberg method, was <0.0001). For visualization purposes, we show the states’ FO across theta phase bins in **Supplementary Figure 4**. Overall, these results indicate that the states are not clearly related to the phase of theta, suggesting they are a different effect from phase precession. **Supplementary Figure 5** shows the HMM states characterisation (i.e., the information reported in **Figure 2**) for each individual animal.

To test whether the HMM states could be influenced by potential shape changes in the data (as seen in hippocampal theta literature ^55^), we run the HMM on simulated data, varying both in shape and frequency. Our results show that the HMM states would be mainly driven by cycle-average frequency changes in a case where the signal changed both in shape and cycle-average frequency, independently (**Supplementary Figure 6**).

To check for spikes bleeding into the LFP that could potentially affect the analysis ^56–58^, we performed a sanity check where we used a sharp low-pass filter at 100 Hz (see **Methods** for more details) to remove the potential effect of spike waveforms on the LFP ^59^. We then run the HMM on the low-pass filtered data and confirmed that the states’ power spectra remained identical to those from the minimally preprocessed data. We compared the HMMs by computing the correlation across the whole recordings between the state time courses obtained from the minimally preprocessed data and those obtained from the low-pass filtered data, across 10 independent HMM runs. We report the correlations, consistently between 0.85 and 0.9 across subjects, in **Supplementary Figure 7**, together with two visual examples of the two correlated state time courses, for different correlation measures. The high correlation values suggested that spike contamination effect did not affect our analyses. We later confirmed this by running the decoding analyses on the low-passed filtered data and comparing with the results on the minimally preprocessed signal.

To investigate whether the state activations indexed the activation of distinct neural subpopulations, we inspected the average firing rate of each neuron during both states. Many neurons showed significant differences in firing rate between the two states, with the HPHT state generally associated with higher firing rates. However, most neurons were active during both states (**Supplementary Figure 8**). To further clarify the relation between firing neurons and state activation, we tried to predict the state activation from the neural activity. We refer to neural population activity as the firing density of all the neurons recorded for each animal, across the duration of a trial, for all the trials performed. Firing density was computed from the binary spike data by means of a Gaussian kernel (see **Methods** for more details). To be able to assess if there is an effect above and beyond the average neural firing rate, we standardised the signal across neurons, for each trial and time point, such that regardless of which state is active the gross amount of firing is the same for all trials. The prediction accuracy for the state’s activation was generally close to chance level (**Supplementary Figure 9**), indicating that there is no clear correspondence between an active state and a specific subpopulation of neurons.

To explore whether the states were related to experimental variables, we examined their temporal distribution relative to trial outcomes (success vs. failure). We found that the states partially reflected trial outcome, with LPLT state being (in three out of four rats) the most active after response for successful trials. For the failed trials, however, each state was on average active for half of the trials (**Supplementary Figure 10**). We also examined the states distributions in relation to relevant task moments, such as odour sampling, response and inter-trial periods. We found consistent patterns of the states’ occurrences in the various moments across subjects, i.e. the LPLT state is the most active during all task-relevant moments, and the distribution of the states is constant during the intertrial time (**Supplementary Figure 11**).

To summarize, our data-driven detected network states showed consistent characteristics across animals, without a one-to-one correspondence between states and behavioural outcome —although the LPLT state was generally the most active during rewarded trials. While the HPHT state typically involved higher firing rate from all neurons, our analyses suggested that the states might not necessarily index distinct subpopulations of neurons.

### Movement does not solely explain state differences or the neural representation of trial outcome

As the experiment involved freely moving animals required to run to the opposite end of the track approximately every fourth trial ^49, 50^, we explicitly tested whether the HMM states and neural representations of trial outcome could be explained by movement. We hence computed the mean and standard deviation of the subjects’ position per state visit across states. The two states did not significantly differ in mean Xp or Yp (the coordinates of the animals along the long and short axis of the track, respectively), but did differ in the standard deviation of Xp and Yp, in three out of four subjects, suggesting that HPHT state visits encompassed a wider range of positions without movement being the primary driver of state assignment (**Table 1**).

**Table 1:** Characterisation of the states by motion. . Average values of the distributions of mean and standard deviation of Xp and Yp coordinates per state visit, and highlighted in green where the distributions are significantly different per Mann-Whitney U test (p<0.0001). The standard deviation distributions on both Xp and Yp coordinates are significant in three out of four subjects, implying the HPHT state covers a wider range of positions on average per state visit.

|  | Mean Xp |  | Std Xp |  | Mean Yp |  | Std Yp |  |
| --- | --- | --- | --- | --- | --- | --- | --- | --- |
|  | LPLT | HPHT | LPLT | HPHT | LPLT | HPHT | LPLT | HPHT |
| Subj 1 | 339.06 | 338.54 | 5.13 | 11.21 | 252.55 | 252.36 | 2.79 | 3.45 |
| Subj 2 | 309.82 | 309.57 | 5.02 | 8.42 | 260.64 | 261.21 | 3.80 | 4.05 |
| Subj 3 | 451.23 | 451.25 | 3.84 | 4.78 | 270.81 | 270.84 | 2.21 | 3.16 |
| Subj 4 | 532.42 | 532.43 | 3.48 | 6.91 | 272.29 | 272.30 | 2.48 | 3.24 |

After checking for locomotion potentially leaking into trial data (see **Methods**), we discarded 13, 2, 18, and 2 trials for subjects 1, 2, 3, and 4 respectively. To test how much movement would explain trial outcome, we used linear classification to predict success and failure from Xp and Yp head position. Head position predicted trial outcome above chance level particularly 1–2 seconds after response - consistent with reward-related behaviour (licking) in successful trials (**Supplementary Figure 12**). To assess how much of the neural signal could be attributed to head motion, we regressed out Xp and Yp position from the spike density data. Decoding accuracy from motion-deconfounded neural activity was slightly reduced relative to the original data (i.e., neural activity not deconfounded for motion), but remained above chance level after response, indicating that head motion is a contributing factor but not the sole driver of the neural signal (**Supplementary Figure 13**) in representing trial outcome. For comparison, decoding accuracy from shuffled neural activity was instead at chance level.

### Theta states reflect neural activity reconfiguration with respect to behaviour

Our tests on the states, movement and neural activity with respect to trial outcome unveiled a complex picture, where the states are linked to trial outcome and movement, but these are not the only drivers of the states assignment. We then asked whether success and failure are represented differently by the neural population activity depending on the theta state. To address this question, we investigated whether the two theta network states influenced how the neural population reflected trial success or failure (i.e. *trial outcome*). Trial outcome may be associated to different cognitive processes, like attentional (which influences the choosing of the correct answer), or emotional (following reward). As before, and for all our subsequent prediction and decoding analyses, the neural population activity across neurons was standardised for each trial and time point - such that regardless of which state is active the gross amount of firing is the same for all trials- and deconfounded for motion (i.e. Xp and Yp position were regressed out from the spike density data).

We used linear regression to predict, one neuron at a time, trial outcome from three factors: the deconfounded neural activity, the states activation and their interaction (the product between deconfounded neural activity and states activation, which is one additional regressor). A schematic of this analysis is in **Figure 1C**. **Figure 3A** shows the effect size of including theta states in the prediction of trial outcome from neural activity (plotted the prediction error averaged across neurons, using the HMM states and randomly sampled states for comparison). Including the theta states in the regression analysis significantly improved trial outcome prediction, reducing the absolute mean error by 10% to 20% in the time points after response (suggesting a link between the LFP spectral properties and reward, instead of attention), with respect to the randomly sampled states. **Figure 3B** shows statistical significance for each neuron, assessed using permutation testing with 10,000 random permutations (p-values corrected for multiple comparisons across time series with the cluster-based spatial-temporal test, see **Methods**). These plots indicate the contribution of state-related regressors (i.e., state activation and state-neuron interaction) to prediction of trial outcome, meaning whether states information improves prediction of success vs failure from neural activity. The plots show that the states’ contributions are relatively consistent across neurons (the clusters of significant time points across neurons). This could hint at state effects manifesting at the level of the whole neural ensemble.

**Figure 3:**
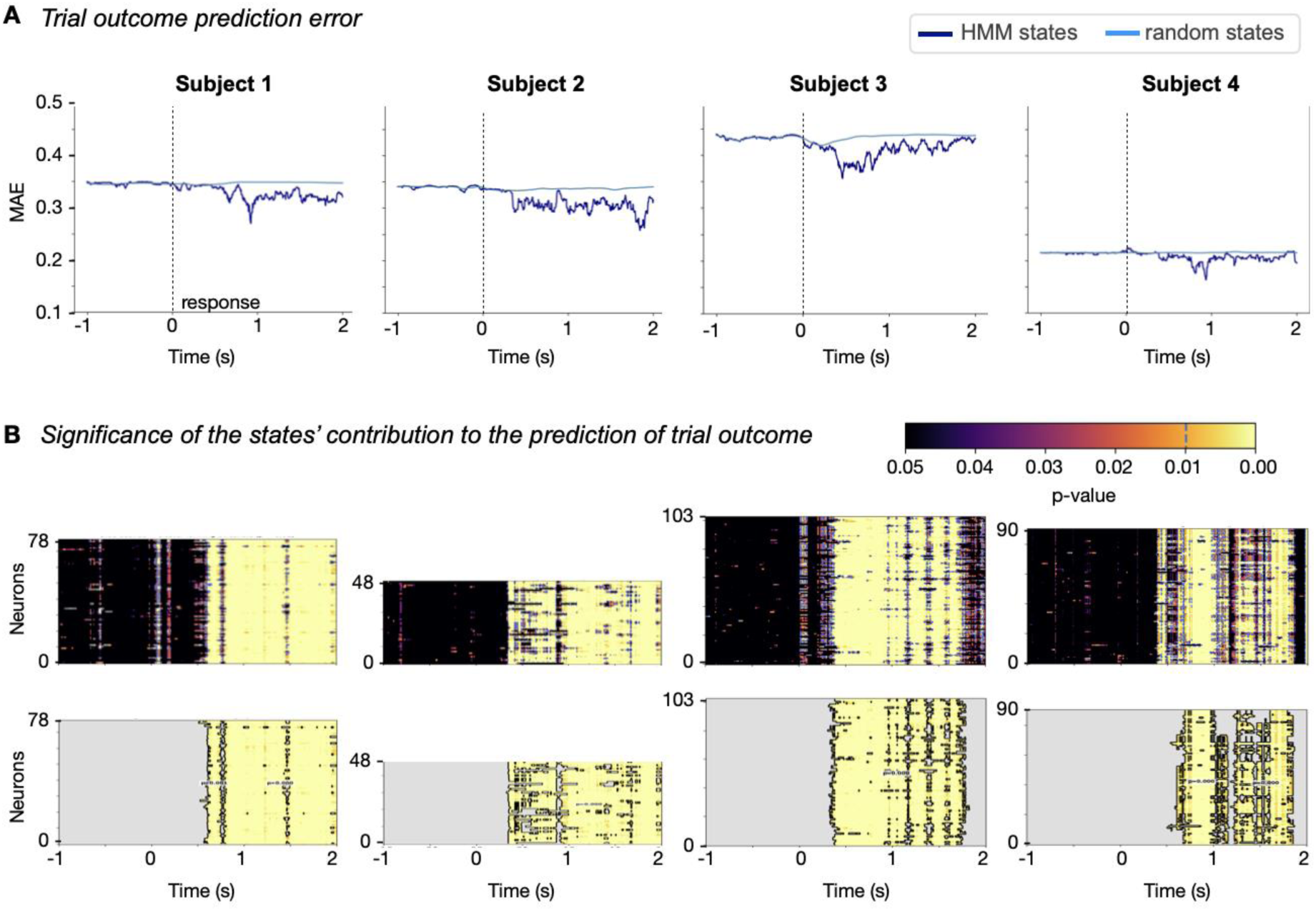
**A.***Trial outcome prediction error*. The plots show, for each subject, the prediction error of trial outcome. The regression analysis was performed per neuron and per time point, including the original HMM states (blue), and then, for comparison, including surrogate states (cyan, 10,000 random permutations). We show the mean absolute error (MAE) of the regression, averaged across neurons (and across permutations). **B.** *Significance of the states’ contribution to the prediction of trial outcome*. P-value of permutation testing analysis, showing how significantly lower is the regression error predicting trial outcome from neural activity, HMM states and their interaction, with respect to randomly assigned states (10,000 permutations). We show the uncorrected p-values (top panel) and spatial-temporal cluster-based corrected (bottom panel), where only the significant clusters with p<0.01 are shown.

To summarize, our interaction analysis showed that accounting for the theta states improves the prediction of trial outcome from neural activity, meaning that the two theta network states reflect changes in how single neurons in a population fire in response to trial outcome, suggesting population-level effects.

### Modulations in mesoscale network activity relate to behaviourally-relevant changes in microscale network activity

Our neuron-state interaction analysis showed that including information on the two network states significantly improved the prediction of trial outcome. We then asked whether the difference between the multivariate patterns of population activity associated with the representation of success and failure is constant across network states. We quantified this difference by using multivariate regression on all neurons at once.

To achieve this, we developed a decoding method that we called *state-conditioned decoding*, where we used the HMM states time courses to separate, per time point, the neural activity into two sets based on the currently active state from the LFP. This separated the trials into two non-overlapping sets per time point, which we then used to decode trial outcome. We quantified differences in population activity patterns between the two states by comparing the decoding accuracy of models trained and tested within the same state (within-state) to that of models trained on one state and tested on the other (cross-state). Figure 1D shows a schematic of this analysis. Our reasoning was that if the two HMM states induced different neural activity patterns, the cross-state decoding would perform worse than within-state. The algorithm enforces that the two per-state decoders use the same number of trials and are balanced in predictive class at each time point (see **Methods**). Of note, we performed the decoding analysis on three out of four rats, as in one of them (Subject 4) the number of failed trials was too low to reliably perform state-conditioned decoding. We noted that within-state decoding accuracy was higher than cross-state decoding for some time points after response in two out of three animals. This difference remained statistically significant after permutation testing (10,000 permutations, cluster-based corrected; see **Methods**) in only one animal. We therefore report decoding accuracies together with their 95% confidence intervals in **Figure 4A**. The trend in one out of three animals supports the interpretation that the two HMM states are associated with partially distinct patterns of neural population activity with respect to trial outcome. We visualized each individual state’s accuracy, and standard accuracy (standard accuracy being the prediction accuracy when decoding from all the trials used in the states decoding), in **Figure 4B**. As observed, the accuracy changes between states and across time. Of note, no one state constantly holds the highest decoding accuracy across time: if one state was always the worst performing, we could reasonably consider whether the states reflected signal to noise ratio distinctions. The fact that the highest decoding accuracy alternates among states across time suggests that the states capture different patterns that become relevant at different time points for the processing of trial outcome. To visually explore the states contributions, we plot the absolute difference in decoders weights in **Figure 4C**: the plot shows that the main differences in the decoders occur for some specific neurons across time and are more pronounced when within-state accuracy is higher than cross-state accuracy (which is also when the two states accuracies differ the most). This plot is only intended as a visualization tool and should be interpreted carefully given that significance was reached in only one out of three subjects.

**Figure 4:**
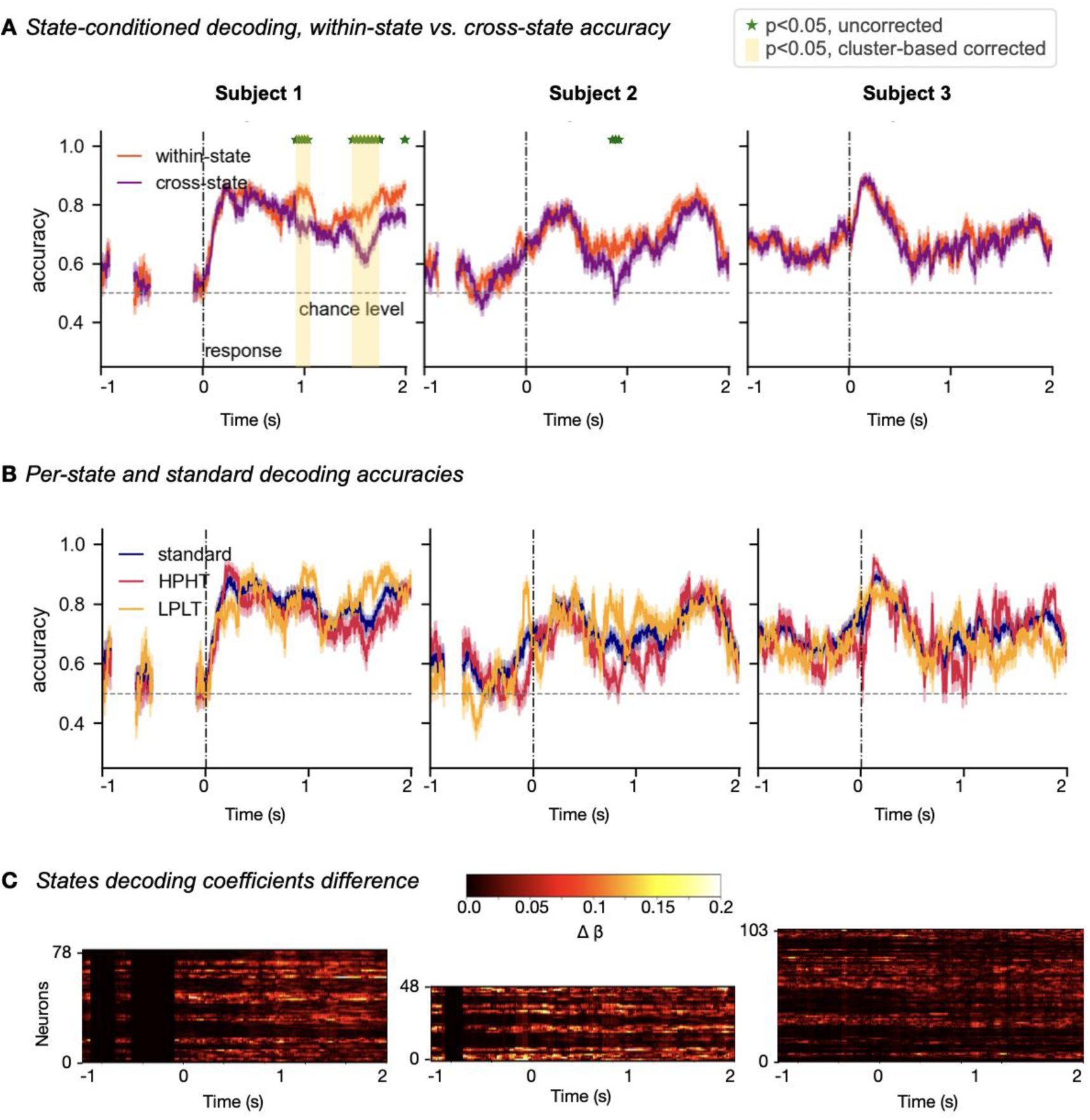
**A.** *State-conditioned decoding, within-state vs. cross-state accuracy*. Comparison between within-state (i.e., training and testing on the same state) and cross-state (training on one state and testing on the other) accuracy, for the state-conditioned decoding analysis, within their 95% CI. For each time point, each state has a different number of trials. If the number of trials for any state at any point is lower than 5 per class, the decoding is not performed (gaps in the plot). The standard decoding is performed with all the trials used for both states, at any timepoint. Permutation testing (10,000 permutations, ensuring class and state-balance at each permutation), is used to assess if the within-state accuracy is significantly higher than the cross-state accuracy, with respect to surrogate states. Green stars indicate which points are significant, p<0.05, uncorrected. Yellow shaded areas indicate which points are significant after cluster-based correction, p<0.05. **B.** Per-state (yellow for LPLT and red for HPHT) and standard decoding (i.e. decoding from all trials, in blue) accuracy, within their 95% CI. States accuracy fluctuate across time points, suggesting the states do not simply capture signal to noise ratio differences in the data. **C.** The absolute difference of the two states decoding weights, for visualization purposes. The main differences occur for most neurons across time, and are most accentuated when the two states have different decoding accuracies.

We visualized the population activity patterns for successful vs failed trials (average across trials) under each state, and showed the absolute difference of these patterns across states (**Supplementary Figure 14**). This plot is intended to complement the visualization of the described effect, that we termed *ensemble pattern reconfiguration*, suggesting that the representation of success vs failure by the entire neural population changes across states.

As a sanity check, we also performed state-conditioned decoding on the HMM states ran on the low-pass filtered LFP signal (see above), confirming that potential spike contamination did not interfere with ensemble pattern reconfiguration effect. We report these decoding results in **Supplementary Figure 15**, not qualitatively different from the decoding results obtained from the broadband states (reported in **Figure 4**).

To summarise, our state-conditioned decoding analysis allowed us to investigate the activity of the neural population per state, so that we could ask whether the neural population differently represented success and failure under the different states. While our results did not reach significance on all the subjects per our permutation testing approach, the effect we found hints at a differential representation of trial outcome for some timepoints after response across the two states. While the neuron-state interaction analysis focused on the states modulation of the firing of individual neurons, state-conditioned decoding analysis addressed changes in the whole population ensemble.

### State-related patterns of trial outcome representation share overlapping characteristics

As observed, even when the within-state decoding accuracy was significantly higher than the cross-state, the latter remained relatively high. Moreover, standard accuracy (i.e., accuracy of the decoding from the neural population without considering the states) was also comparable to that of within-state decoding. This is due to similarities between the two states’ neural ensemble activity patterns. If the patterns were entirely distinct, cross-state accuracy would be at chance level, and standard accuracy would also be poor; while if they were nearly identical, cross-state accuracy would match that of within-state (and standard accuracy would be higher than both). To explore this effect, we simulated datasets with varying degrees of overlap between two data subsets, from fully similar, to fully distinct (see **Methods**). The simulations indicated that our experimental data was best matched when the subsets had an information overlap between 70% and 80% (**Supplementary Figure 16**). This suggests that, while the two states are distinct, they still share substantial amount of overlapping information.

## Discussion

This study explores a form of oscillation-to-neuron relation in the hippocampus of rats performing an odour-memory, non-spatial task, which we refer to as *ensemble pattern reconfiguration*. Using an unsupervised approach, we identified two distinct theta states, low-power-lower theta (LPLT) and high-power-higher theta (HPHT). Using decoding analyses, we showed that the states were linked to differential neural ensemble activity reflective of trial outcome, meaning that the difference in neural population representing success vs failure might change across states. These states did not appear to recruit different neurons but rather were linked to changes in how the same population processed information. This suggests a mechanism of functional multiplexing, where the same neurons represent stimuli differently depending on the network configuration ^14,15,20^.

Previous research on oscillation-to-neuron modulations in the hippocampus has primarily focused on phase precession, a phenomenon where the phase of theta oscillations influences the firing of single neurons during navigation ^48^. Our work uncovers a different type of link between neural population activity and oscillations, where differences in neural firing patterns representing trial outcome are reflected in changes in amplitude and frequency of oscillations within the theta band.

Our findings link the activity of single neurons and oscillations at the network level, contributing to the literature that aims at bridging different spatial scales in the brain. Our conclusions are consistent with the concept of “frequency sliding” in the LFP data ^43^, i.e. frequency of oscillations not constant in time, but “sliding” within a small range during cognitive processes and resting state, affecting the spiking threshold of single neurons. In Cohen (2014), the author speculated that frequency sliding could serve functional purposes, as the slower frequencies could induce faster but more variable spiking time, and the faster frequencies would instead trigger slower but more precise firing. In this view, our work might help shed light in the functional mechanisms used for information encoding in the brain.

Even though the HMM that we used to detect network states is a purely data-driven model with no neurobiological assumptions or knowledge of the task, the two distinct state peaks within the theta band are reminiscent of the two previously identified hippocampal theta modes, generated from regions projecting to the hippocampus, and related to different behavioural scenarios: Type I (∼7–12 Hz, atropine-resistant) during locomotion and Type II (∼4–7 Hz, atropine-sensitive) during attentive immobility ^60–69^. In awake immobility, theta has been observed to transiently accelerate toward the upper band yet remaining pharmacologically of Type II (that is, sensitive to atropine) ^70^. The two data-driven states we observed here show some relation to movement in the experiment (namely, one state covered a wider range of positions), consistent with the traditional distinction of the two types of theta modes in the hippocampus in literature. The states, however, do not seem to be solely driven by motion, as they are both involved during the trials, performed in a constant spatial location, suggesting that coordinated shifts in theta frequency and power might be associated with ensemble-level reconfiguration in reflecting trial outcome. The exact biological function of these network states remains speculative. Mechanistic inference would require interventions to pinpoint the pathways driving the ensemble modulation, coupled with controlled task manipulations to establish functional relevance. Theta frequency also tracks internal states across behavioural contexts: motor preparation ^71^, environmental novelty ^72^, and higher arousal states such as risk-taking ^73^, anxiety ^74^, and social interaction ^75^. We acknowledge that residual motion contributions cannot be fully ruled out. Future work with head-fixed preparations would allow a cleaner dissociation between cognitive and motor contributions. Further investigation would also be needed to explore how these states relate to cognitive processes such as task engagement, decision making or learning ^47^.

Importantly, we do not claim the existence of exactly two theta states in the hippocampus. This number of states is simply the level of granularity afforded by the current data, but is sufficient to show the co-occurrence between the spectral properties of theta and the distributional properties of the neural firing with respect to behavioural context. This does not exclude that the effect we have uncovered is supported by variations within a continuous distribution, rather than by categorical distinct states. In any case, it still reveals a biologically relevant function in that changes within power and frequency of oscillatory patterns are linked to nuanced changes in population activity in its relation to behaviourally meaningful variables. Also importantly, whereas we are here describing an oscillation-to-neuron modulation, we do not claim an underlying unidirectional causal flow from one to the other: oscillations and neural firing are two properties of the same system that emerge together.

Our focus on the theta band was dictated by its known relevance in the hippocampus, and because it is prominent in the data. However, the data showed oscillations across the broadband frequency spectrum, particularly in both beta and gamma bands. This is consistent with research on odour-related tasks (typically in the beta band ^76^) and on temporal processing (typically associated to gamma ^77^). It is important to note that, since we have not identified frequency and power as independent features of our states, we cannot disentangle them, implying we are not able with this analysis to further describe whether either one or both features are the main driving characteristics of this effect.

We acknowledge the small number of animals analysed here as a limitation in this study. Future research may explore whether these findings generalize to other animals, brain regions, or tasks.

In conclusion, this study suggests that changes in the network state, identified here as subtle changes in oscillatory power and frequency, are correlated with how the neural population activity represents behavioural information, thus challenging static models of neural coding ^20^. That is, the hippocampal circuit might process complex variables such as success and failure in multiple ways depending on the state of the system. This underscores the role of theta oscillations in the hippocampus, reflecting a flexible neural information processing landscape, and possibly contributing to the brain’s adaptation to changing cognitive demands ^6, 78^.

## Methods

### Dataset

The data were collected by Shahbaba and colleagues ^49, 50^ and made publicly available. The dataset consists of local field potential (LFP) and spike recordings from the dorsal CA1 region of five rats performing an odour-memory task. Rats learned a sequence of five different odours. In the experiment, they were presented with an odour at a time and had to identify it as either “in sequence”, if it was naturally following the learnt sequence, or “out of sequence”, if it was placed in a wrong position within the learnt sequence. Rats would judge the odour in sequence by holding their nose in the response port for at least 1.2 s to receive a water reward, or out of sequence by withdrawing their nose before 1.0 s to receive the reward. The data were recorded when rats had learnt to perform the task very well, so that success rate is above 80%. For each rat, one session was recorded. Across animals, the number of trials varied between 140 and 300 and the number of neurons detected varied between 40 and 80. The original sampling frequency of the data was 1kHz. While the data of four out of five rats exhibited similar spectral characteristics, the power spectrum of the fifth rat was considerably different (see **Supplementary Figure 17**). This important difference, possibly due to a difference in electrodes placement (see Allen et al., 2016 ^50^ for details in the electrode placement procedure), induced us to exclude subject 5 from our main analyses for consistency. For the sake of completeness, however, all analyses were performed also on Subject 5, reported in **Supplementary Figure 18**.

### Data preprocessing

The LFP channels recorded from the animals’ hippocampus varied between 18 and 24 across animals. Activity was very highly correlated across electrodes. On the raw LFP data per subject **Z**, with dimensions (number of timepoints by number of channels), we applied principal component analysis (PCA; ^79^) to obtain **Z,** with dimensions (number of timepoints by number of principal components). We then kept only the first principal component **z**, with shape (number of timepoints by one), which explained between 60% to 75% of the variance in each subject (subject 1: 65%, subject 2: 70%, subject 3: 60%, subject 4: 63%, subject 5: 75%). To remove noisy artifacts at lower frequencies, we high-pass filtered **z**, with a cut-off frequency of 4Hz. Finally, **z** was downsampled to 250Hz. To control for potential spike-LFP bleed through effects, we designed a sharp low-pass digital filter with a Kaiser window ^80^ to attenuate by 100% the power on the LFP signal above 100 Hz, within a stopband of 1 Hz. The low-pass filtered data **z_<100_** was then used for a parallel analysis, and results were compared with the minimally preprocessed data **z**.

We computed the spike density **X** from the raw spike data by means of a Gaussian kernel ^81^ (10ms width). This way, the spike data (originally a binary vector per neuron) was represented in a continuous form for subsequent analyses: **X**, with shape (number of trials by number of neurons, by timepoints per trial) per subject. The data were then standardised per trial and per timepoint, across neurons, so that the population activity had the same mean and standard deviation for each trial and for each time point. This way, we maintained the relationship between the neurons, but we rescaled the overall activity at each timepoint and each trial. We did this so that the neurons activity was not trivially linked to the states activation (since higher average firing rate occurs when the state HPHT is active). As with the LFP, the spike densities were downsampled to 250 Hz.

### Motion analysis and deconfounding

As the experiment involved freely moving animals, we explicitly tested whether the HMM states and the neural representation of trial outcome could be explained by movement-related confounds. We distinguished between two types of movement: locomotion (full-body translation along the track) and head motion (local movements at the port, including head turns and postural adjustments), and addressed each separately.

### Movement data

The Xp and Yp, coordinates of the animal head’s position along the track - recorded at a 30 Hz sampling rate – were upsampled to the 250 Hz of the neural data, and smoothed with a gaussian kernel of width of 500ms to attenuate noise. The Xp dimension corresponds to the long axis of the track, 150 cm long, while the Yp dimension corresponds to the short axis, 9 cm wide. These coordinates are expressed in the reference system of the tracking probe, where the coordinate [0,0] is outside the track, with the Xp track between coordinates 50 to 1000 (odour and reward port at approximately Xp = 60–120 and the reward port at opposite end of the track at approximately Xp = 900-1000), and with Yp coordinates spanning between 100 and 500. Note that the exact scale in physical units (cm) should be deduced according to the calibration used in the experiment.

### Characterisation of state-related movement

For each state visit (a state visit being a collection of consecutive time points assigned to one state), we computed the mean and standard deviation of the Xp and Yp position of the animal. We tested for significant differences in these distributions across states using the Mann-Whitney U test ^82^.

### Locomotion exclusion

No universally accepted definition of locomotion exists for freely moving rodent recording experiments, with thresholds varying substantially depending on task structure ^65,83,84^. We therefore defined an operational criterion adapted to the spatial constraints of our task. We classified a trial as locomotion-affected if the animal displaced more than approximately one body length ( ∼20 cm) along the long axis of the track (Xp dimension) within any 500 ms window falling within the extended trial epoch of −1.0 to +2.5 s around response. This criterion corresponds to a minimum translational speed of ∼40 cm/s along the track axis, and it excludes only clear running bouts, not local head or postural movements. Applying this criterion, we excluded 13, 2, 18, and 2 trials for subjects 1, 2, 3, and 4 respectively.

### Head motion deconfounding

After discarding locomotion-affected trials, we quantified how much residual head motion (Xp and Yp position) could predict trial outcome. To do so, we cut the position data into trials, so to have data **M** with dimensions number of trials by two (Xp and Yp dimension) by time points. We then standardized **M** across time per position dimension, and finally used class-balanced linear classification (ridge classifier, L2 penalty = 0.001) to predict trial outcome from **M**. We then treated head motion as a confound in all neural analyses by regressing out **M** from the spike density data **X** (per neuron and per time point) using linear regression. The residuals of this regression **X_d_** were used in place of the original spike density for all subsequent motion-deconfounded analyses.

### Definition of states through the HMM-TDE

To detect network states, we used a time-delay embedded hidden Markov model (HMM-TDE; ^53^) on the preprocessed LFP data (see above). The HMM is a data-driven, unsupervised method that automatically detects recurring patterns of activity in the data, referred to as *states*. Importantly, being data-driven, the HMM estimates the states agnostic of any experimental paradigm or trial information in the data. What defines a state depends on the choice of *observation model* of the HMM. In this case, the observation model involves time-delay embeddings (TDE) of the data, focusing on the autocovariance of the signal. Previous work from the authors investigated the sensitivity of the HMM-TDE to various signal characteristics, showing that on one-channel data, the HMM-TDE reliably captures the spectral properties of the signal ^54^. The HMM gives a representation of the time-series data in terms of probabilistic sequences of states. From the data, the inference estimates: the states’ autocovariance, the probability of transitioning to a different state or of remaining in the same state, and the initial probabilities. The HMM also outputs the probability of each data point to pertain to a state. An additional algorithmic procedure returns the Viterbi path ^85^, a categorical state assignment (instead of a probability) for each time point. Both the probabilities and the Viterbi path are referred to as states time courses.

The user-defined hyperparameters of HMM-TDE are:

- the number of states K;
- the lags L: defining the length of the window in which to estimate the data autocovariance (from −L to +L around each time point), which affects the spectral scales that the states can capture;
- the parameterδmanipulating the prior probability of remaining in the same state, which influences the states switching rate.

Here, because of the high class-imbalance of the data with respect to the decoding variable (i.e., data had few failed trials), we chose only *K*=2 states. The lags *L* and the prior probability parameter δ were chosen such that the states would dwell for at least 500ms on average, forcing the states switching rate to be slow enough, so to be able to capture cognitive or physiological processes (typically, of the order of a second ^86^). In this specific case, we chose *L*=7. δ=1bn for rats Buchanan, Stella and Barat, and δ=10m for Mitt. See Masaracchia *et al*. (2023) ^54^ for the rationale on the effect of modulating the hyperparameters on HMM-TDE. To characterise the states, we computed their power spectrum, lifetime, and fractional occupancy.

We explicitly examined the HMM-TDE ability to capture finer details in frequency. We used the same pipeline as in Masaracchia et al. (2023) ^54^ to simulate data varying only in frequency between 7 and 9 Hz, and run the HMM-TDE with the same settings as in this experiment (i.e., K=2, L=7, δ= 1bn). We characterised the states by computing their power spectrum.

The HMM-TDE model, as well as the functions to estimate the states characteristics and power spectra, was implemented and made publicly available within the python GLHMM toolbox on GitHub: https://github.com/vidaurre/glhmm ^87^.

We then quantified statistically the difference between power and frequency of the two states. To do so, we computed the power spectrum of each state occurrence that was longer than 50 timepoints (minimum number of data points to compute a power spectrum, arbitrarily chosen). We extracted the maximum power and its corresponding frequency for each valid (i.e., longer than 50 timepoints) state visit, obtaining a distribution of power and frequency for each state. We finally computed whether the difference in the power and frequency distribution between states were significant by means of the Mann-Whitney U test ^82^, and corrected for multiple comparisons using the Benjamini/Hochberg method ^88^.

To examine the states relation to theta phase, we computed the periodicity of occurrence of the states. This was done by computing the power spectrum of the state time courses, i.e. the probability of each state to be active at any time point. This measure shows the frequency of activation of the states (and because the states are exclusive, they have the same activation frequency). To explicitly link theta phase to the state time courses, we computed the theta phase of each state visit and compared the two states distribution in angles. To do so, we first extracted the phase from the LFP signal, band-pass filtered in theta band (4-12 Hz). We then grouped the theta phase time course into two groups, according to which state was active at each time point. This gave us two distributions of angles, one for each state, that we compared by means of the Watson U^2^ test ^89^. This test is similar to the Mann-Whitney test but for circular data. To visualize the states distribution across theta phase, we divided the time courses in bins of 1/5^th^ π angle each, and computed the fractional occupancy (FO) of the states for each bin (**Supplementary Figure 4**).

### Simulating oscillations changing in shape

To test whether the HMM states could be driven by changes in the oscillation shape, we generated synthetic oscillatory signals using a process based on *Genephys*, a generative model of electrophysiological data (publicly available at https://github.com/vidaurre/genephys ^90^), in which frequency *f*, waveform shape modulation *m*, and amplitude *a* evolve as independent autoregressive processes of order one around prescribed targets. This structure introduces slow stochastic wander while maintaining temporal continuity to reproduce the variability observed in neural recordings as follows:

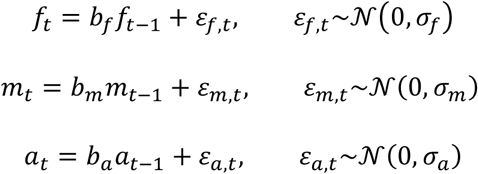

Wheref *b_f_, b_m_* and *b_a_* are the autoregressive weights of the process, and the various *ε* are noise contributions, normally distributed with zero mean and corresponding standard deviation *σ*. To change the shape, we modulated the phase of the sinusoid *φ* according to a nonlinear cumulative update, with the carrier frequency determining the period and the modulation parameter m distorting cycle symmetry:

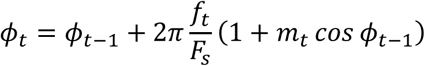

Here *F* _*s*_ is the signal sampling rate. Thus, when *m* = 0, the signal reduces to a sinusoid. Non-zero values of *m* accelerate phase progression during one half-cycle and decelerate it during the other, producing asymmetric waveforms without altering the period.

The observed signal was obtained by modulating a sinusoid with these processes and adding Gaussian measurement noise:

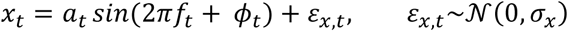

To build continuous time series, we concatenated segments with randomized durations with a minimum bound of 10 seconds and rescaled to a fixed total duration of 5 minutes, ensuring heterogeneous segment lengths. We then specified frequency variation (7 - 9 Hz), shape variation with *m* ∈ [0, 1], combining both independently across time. This design allowed us to control for frequency shifts, shape modulations, and their interaction on downstream analyses.

### Predicting state activations from neural activity

To check whether the states captured the activity of different subpopulations of neurons, we used supervised classification to predict the state activations from the neural activity (deconfounded for motion) **X_d_**. As the HPHT state generally involved higher firing rate for most neurons, the spike density was standardised per trial and per time point, across neurons. This way, we were able to capture differences that were above and beyond gross levels of activation. The decoding algorithm used was a ridge classifier with a fixed L2 penalty of 0.001. Because of the imbalanced distribution of the states across trials and time points, we implemented a class-balanced training-test decoding analysis: we bootstrapped sets with the same number of trials for train and testing for each state, per time point. We repeated the procedure 100 times, randomly sampling the trials involved in the analysis for each repetition. This way, we ensured a consistent class balance for all time points. We measured accuracy as an average across all bootstrap repetitions, per time point.

### Predicting trial outcome from neural activity and state activations

To determine the contribution of network states to the prediction of trial outcome, we used standard ridge regression, and permutation testing. Specifically, we aimed at predicting the trial outcome variable Y from three factors: the neural activity (deconfounded for motion) **X_d_**, the states activation and their interaction (i.e., the multiplication between the two terms). For each neuron and each timepoint, we have the following regression equation:

where ***X_d_*** is the value of the kernel-smoothed activity deconfounded for motion, for each neuron and for each timepoint, a vector of length number of trials, G indicates whether state HPHT or LPLT is active (a vector of length number of trials, with elements equal either to 1 or −1), *w* are the regression coefficients (one number for each *w*), and the intercept *b* (a vector of length number of trials, with one value, repeated for each trial) is related to the class balance. In particular, the *w*_*XG*_ coefficients reflected the interaction between the states activation and the neurons activity. We computed the regression coefficients that minimize the prediction error, i.e., the mean absolute error between the predicted and true value of Y. We used a small L2 penalty of 0.001. Here, the rationale for using Ridge regression instead of logistic regression is that for a binary classification problem the least square error metric would allow for a more nuanced description of the prediction output ^91^.

To quantify the significance of the states’ contributions in explaining trial outcome, we used permutation testing. Specifically, we compared the mean absolute error from the original regression model, with the network states identified by the HMM, against models using randomly shuffled states (10,000 random permutations). The shuffling was performed by swapping the probability of each state in randomly selected 50% of trials (where swapping 100% of the trials in this case corresponds to simply swap the states labels), for each permutation. This shuffling procedure ensured that only the states’ effect (and not the overall prediction power) was tested. This gave a p-value for each neuron and for each time point, that we corrected for multiple comparisons using a cluster-based spatial-temporal correction^92^.

The correction proceeded as follows. For each neuron and time point, we defined an effect score as the difference between the mean null MAE (averaged across permutations) and the observed MAE: a positive score indicates that the observed prediction error is lower than expected under the null, i.e. that the model genuinely outperforms chance. We computed a one-tailed p-value at each neuron–time-point as the fraction of permutations for which the null MAE was less than or equal to the observed MAE; a small p-value therefore indicates that the observed prediction was better than nearly all permutation runs. Points with p < 0.05 were thresholded to form a binary map, and contiguous regions in the neuron–time plane (allowing adjacency both along the time axis and across neurons) were labelled as clusters. The mass of each cluster was defined as the sum of the effect scores across all neuron–timepoint pairs within the cluster; this weights the cluster statistic by effect strength rather than cluster size alone. To build the null distribution of maximum cluster masses, the same procedure was applied to each of the 10,000 permutation runs: the permuted p-values were thresholded at p < 0.05, clusters were identified in the permuted data, and the maximum cluster mass was recorded. A cluster in the observed data was declared significant if its mass exceeded the 95th percentile of this null distribution (cluster-level α = 0.05).

### State-conditioned decoding

To investigate the effect of the mesoscale activity (represented by the two spectrally-specific network states) on the patterns of population activity reflective of success and failure, we developed a method to decode trial outcome from neural activity (deconfounded for motion). To be able to predict trial outcome from the population activity across states, all network states needed to be involved (at least to some extent) in both successful and failed trials. However, given the skewed nature of trial outcome (meaning, rats performed successfully more than 80% of the trials), this condition could only be fulfilled by using the minimum number of two network states. Note that the number of states is a matter of granularity in the analysis, and does not mean a biological claim. We first used the HMM state time courses (indicating which state was active at any given point) to divide, at each time point, the neural activity into two non-overlapping sets of trials, each related to the current active state. We then used a ridge classifier (with a small L2 penalty of 0.001) to decode trial outcome from each of the two state-related sets independently, for each time point.

Due to the high class-imbalance of the data, we implemented a custom class-balanced training-test procedure. The procedure consisted of using only a subset of data for training and testing, the same amount from each class. We repeated this step 200 times, randomly sampling the trials for training and testing from each class at each repetition. This gave us a measure of accuracy and a confidence interval for each state and each time point. For example, if at time point *t* the LPLT state-related set contained 100 successful and only 20 failed trials, we would run the LPLT-related decoder at time point t by using only 16 out of the 20 failed trials for training and 4 trials for testing, and 16 trials for training and 4 for testing from the successful trials pool; we would randomly sample (with no repetition) the trials to use for training or testing for each of the 200 permutations. To ensure comparability of decoder performance and weights across the two states, we additionally enforced state balance across decoders: at each time point, the two per-state decoders were constrained to use the same number of trials. When one state had more trials available at a given time point, a random subset matching the size of the smaller state was drawn at each bootstrap repetition. This ensures that any observed differences in decoding accuracy between states are not driven by differences in trial counts. We called our method *state-conditioned decoding*.

To quantify the difference between the two network states-related patterns of activity reflective of trial outcome, we compared the decoding accuracy of models trained and tested within the same state (*within-state*) and the accuracy of models trained on one state and tested on the other (*cross-state*). In all cases the models were tested on unseen data during training.

To assess the significance of the difference between the state-related population activity patterns, we performed permutation testing. This consisted of randomly assigning trials at each time point to one or the other state, while respecting the original state- and class-balance, for 10,000 random permutations. For each permutation, we performed state-conditioned decoding on the random sets of trials (enforcing a class-balance and a state-balance at each permutation) and computed within-state and cross-state decoding accuracy.

The difference between within-state and cross-state decoding accuracy in randomly assigned states was compared to the original difference (i.e., in the analysis performed with the original HMM states), for each time point. We assessed the significance of this difference (within minus cross-state decoding accuracy) in the HMM versus the random states case, by computing a p-value and correcting for multiple comparisons using a cluster-based permutation approach^92^. Specifically, at each time point we computed the observed difference between within-state and cross-state decoding accuracy under the original HMM state assignments. A time point was considered above threshold if the observed difference exceeded the 95th percentile of the pointwise permutation null distribution (one-tailed, α = 0.05). Clusters were defined as contiguous runs of above-threshold time points. For each cluster, a cluster-level statistic was computed as the sum of the observed differences across all time points in the cluster. To assess the significance of each cluster, we built a null distribution of maximum cluster statistics: for each of the 10,000 permutations, we applied the same thresholding procedure to the permuted differences and recorded the maximum cluster statistic. A cluster was declared significant if its statistic exceeded the 95th percentile of this null distribution (cluster-level α = 0.05).

### Simulations

In order to better understand the results from our state-conditioned decoding, especially why the cross-state decoding accuracy was relatively high, even when significantly lower than the within-state decoding accuracy, we used simulations. We generated two sets of data with varying degrees of similarities, run state-conditioned decoding on the synthetic datasets, and compared the results from the various scenarios to our experimental results. For comparison, we also run standard decoding, i.e. decoding without distinguishing between one or the other set of data.

For this analysis, we generated two random sets of points and the encoding weights that would make the data carry information. The various scenarios, depicting various degrees of similarities between the two datasets, were controlled by the encoding weights: the two sets would carry identical information (and simply be part of one bigger dataset) if the encoding weights were identical; the scenario with completely different encoding weights represented the extreme of non-overlapping information between the two sets. Every other scenario in between was generated by using overlapping weights, controlled by the parameter α. Here follow the equations of our generative model:

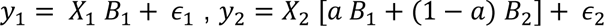

Where *X*_1_ and *X*_2_ are random data generated as ∼*N*(0,4); the dimensions of *X*_1_ and *X*_2_ were made comparable to our experimental data, so we used N=100 as number of trials and p=50 as number of neurons. *ϵ* _1_ and *ϵ* _2_ are random gaussian noise distributed as ∼ *N* (0,1); *B* _1_ and *B* _2_ are the encoding weights both distributed as ∼ *N* (0,4); and *a*, ranging between 0 and 1, indicates the degree of similarity between the two sets of data.

Then, we run the decoding analyses to predict the encoded information (and the encoding weights). Within-set and cross-set decoding were run on the two synthetic sets of data, to decode *y*_1_ and *y*_2_. Standard decoding was run on the concatenated sets to predict an overall y (*y*_1_ and *y*_2_, concatenated). The decoding accuracies obtained were then compared to the experimental results.

## Data & code availability

The data used in this study were pre-collected by Norbert Fortin’s lab and made publicly available (see ^49^). The code used to perform all the analyses is available on GitHub at https://github.com/LauraMasaracchia/state-conditioned_decoding.

## Conflict of Interest

The authors declare no competing financial interests

## Acknowledgements

DV is supported by a Novo Nordisk Foundation Emerging Investigator Fellowship (NNF19OC-0054895) and an ERC Starting Grant (ERC-StG-2019-850404). The authors would also like to thank the Wellcome Trust for support (106183/Z/14/Z, 215573/Z/19/Z). Finally, the authors would like to thank Norbert Fortin and Keiland Cooper for providing the raw data, together with task information and insights into the data structure.

## SUPPLEMENTARY FIGURES

**Supplementary Figure 1:**
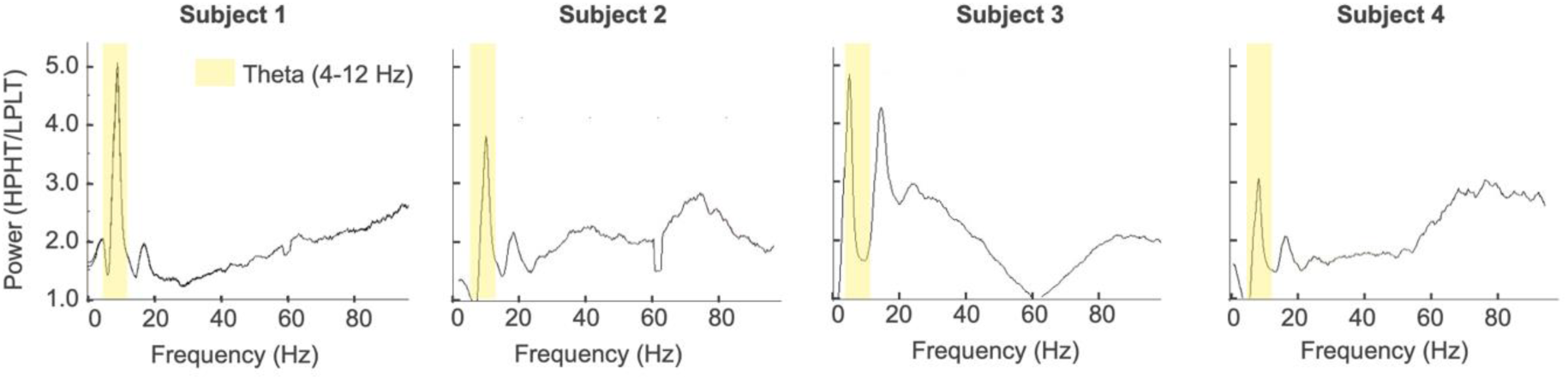
Power difference between HPHT and LPLT states. Here plotted the ratio between the two states power spectrum (HPHT / LPLT), to better visualise the difference in power between the two states proportionally to the overall broadband power spectrum.

**Supplementary Figure 2:**
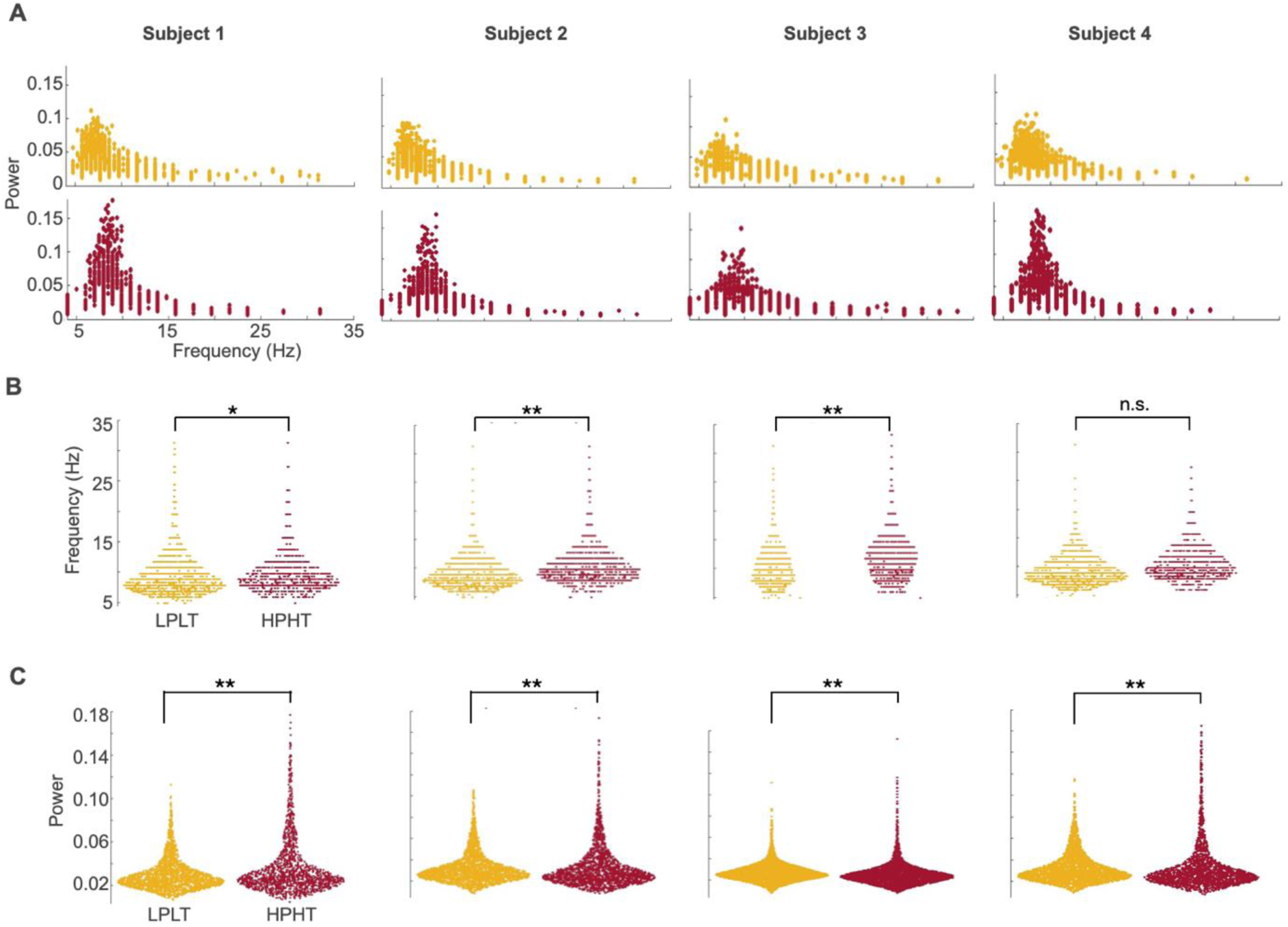
Statistical characterization of the states. **A** *Power spectrum distribution of the states.* For each state visit longer than 50 timepoints, the max power and its corresponding frequency is plotted, for each state, for each subject. **B** *Frequency distribution of state visits.* The frequency of each state visit longer than 50 points is plotted as a distribution, per state and per subject. **C** *Power distribution of state visits.* Like in **B**, but for the max power of each state visit counted. The two state distributions (frequency distributions in **B** and power distributions in **C**) are then statistically compared by means of the Mann-Whitney U test (for each subject). Here represented with * a p value <0.05, and with ** a p value < 0.0001.

**Supplementary Figure 3:**
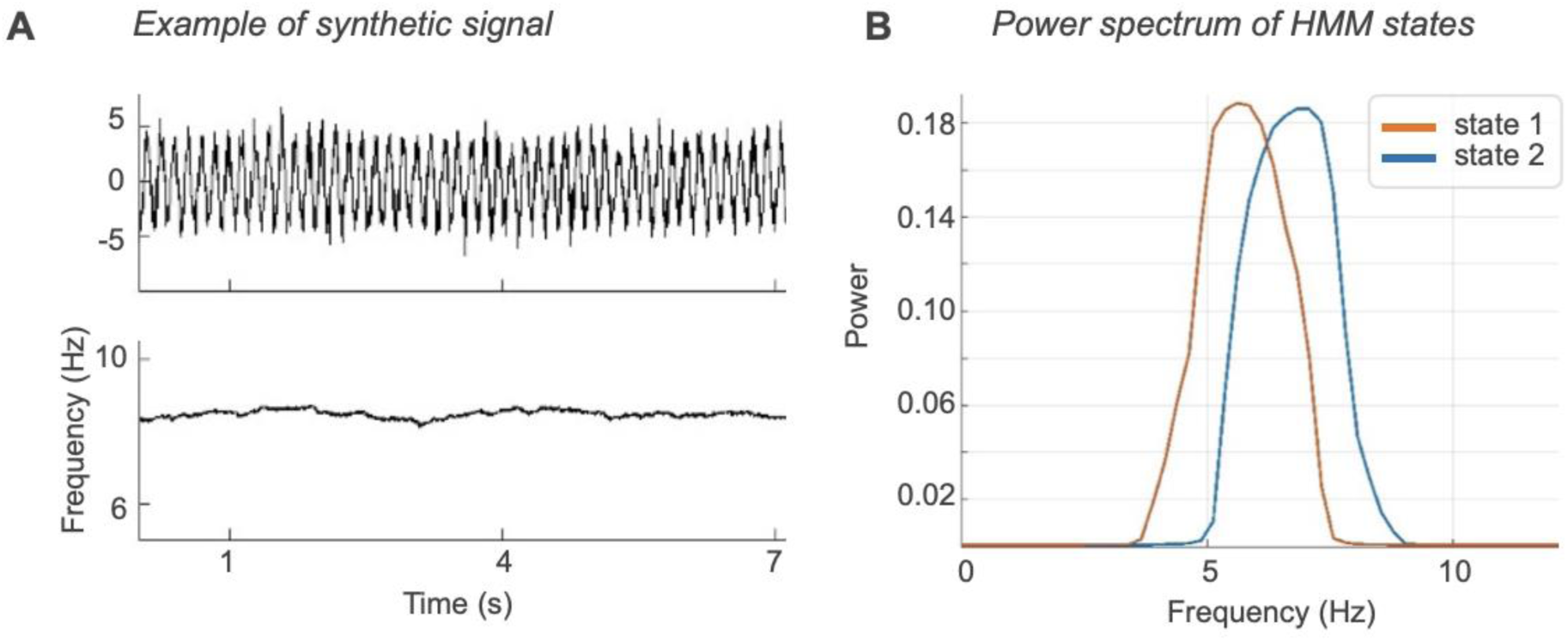
Testing HMM sensitivity to frequency. **A**. *Example of synthetic signal* used to test the HMM sensitivity on frequency (top panel), together with its instantaneous frequency (bottom panel). The signal is simulated as a simple sinusoid with amplitude a= 5, Gaussian noise with standard deviation = 1, and instantaneous frequency determined as a semi random walk bounded between 7 and 9 Hz. **B.** *Power spectrum of HMM states*. Here, the HMM settings are identical to those in our experiment: K=2, *L*=7. δ =1bn. The power spectrum of the states shows a peak frequency difference of about 1Hz.

**Supplementary Figure 4:**
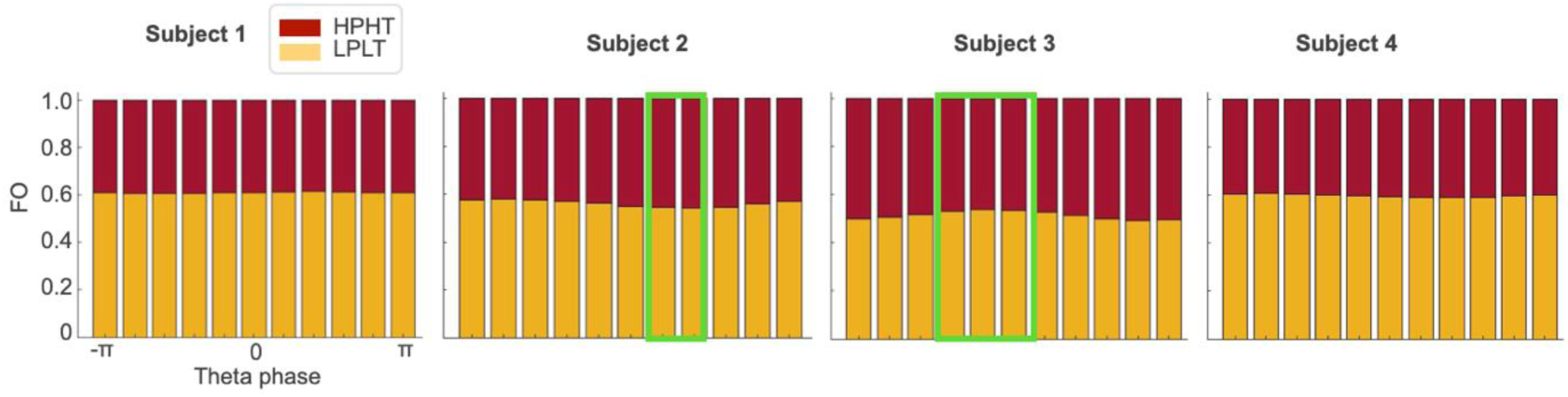
States fractional occupancy (FO) as a function of theta phase. For each theta phase bin of 1/5 π, the states FO is computed and plotted cumulatively. The green window highlights significant changes within the distribution of states FO across phase. This serves as a visual representation of the states distribution around theta phase. For the details on the analysis, see **Methods**.

**Supplementary Figure 5:**
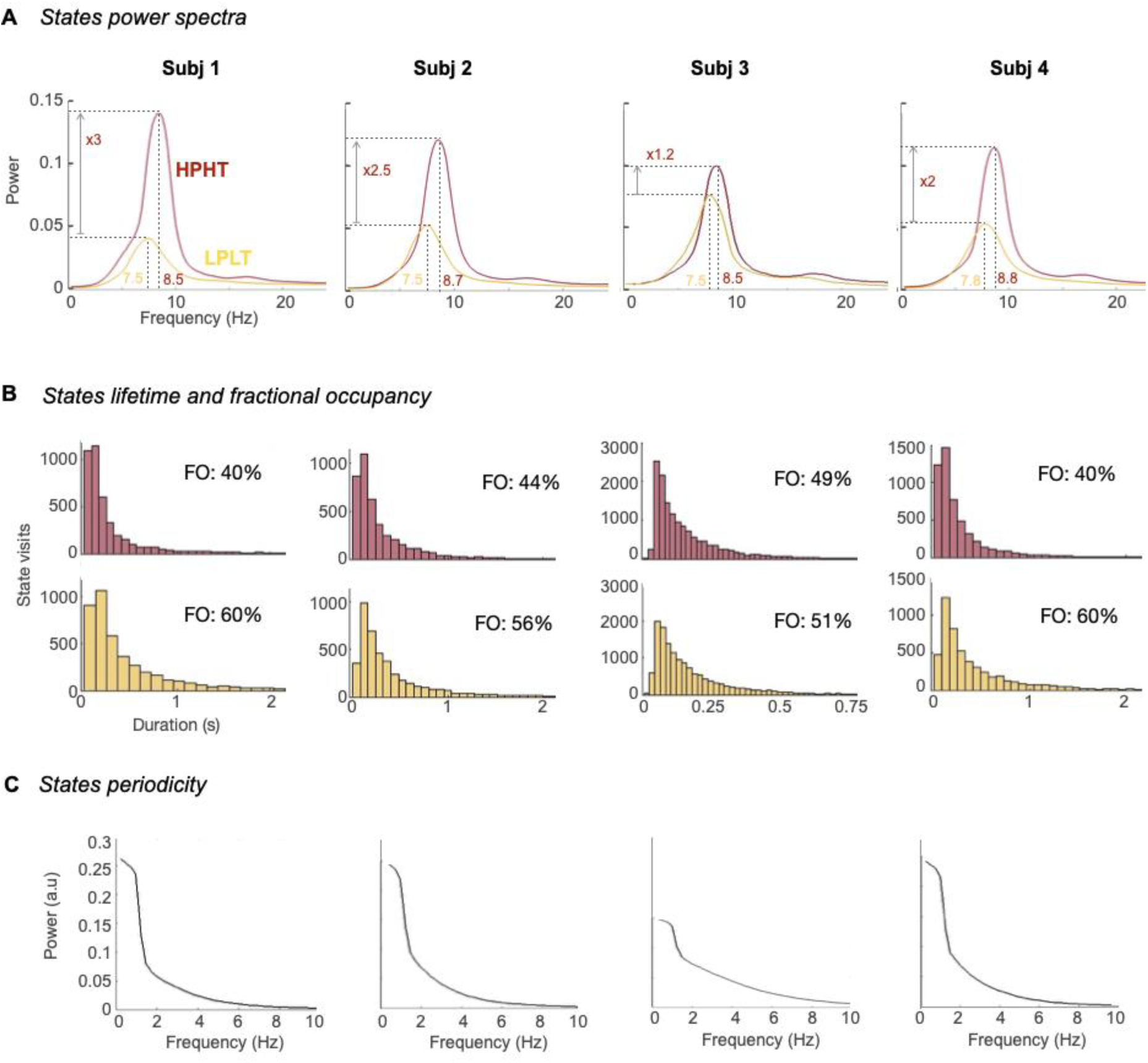
States characterization per subject. **A.** *States power spectra.*Frequency and power content of each state for each subject individually. The HMM consistently finds a high-power, higher theta (HPHT, red) and low-power, lower theta (LPLT, yellow) state. **B.** *States lifetime and fractional occupancy*. The count of the time length of each state occurrence, per state and per subject. Fractional occupancy (FO) accounts for the overall presence of a state in the entire dataset. Both FO and states lifetime are comparable across states. **C.** *States periodicity.* The period of each state activation, per subject. States reoccur typically with a period around 1-2 Hz, meaning that each state occurs on average once to twice per second.

**Supplementary Figure 6:**
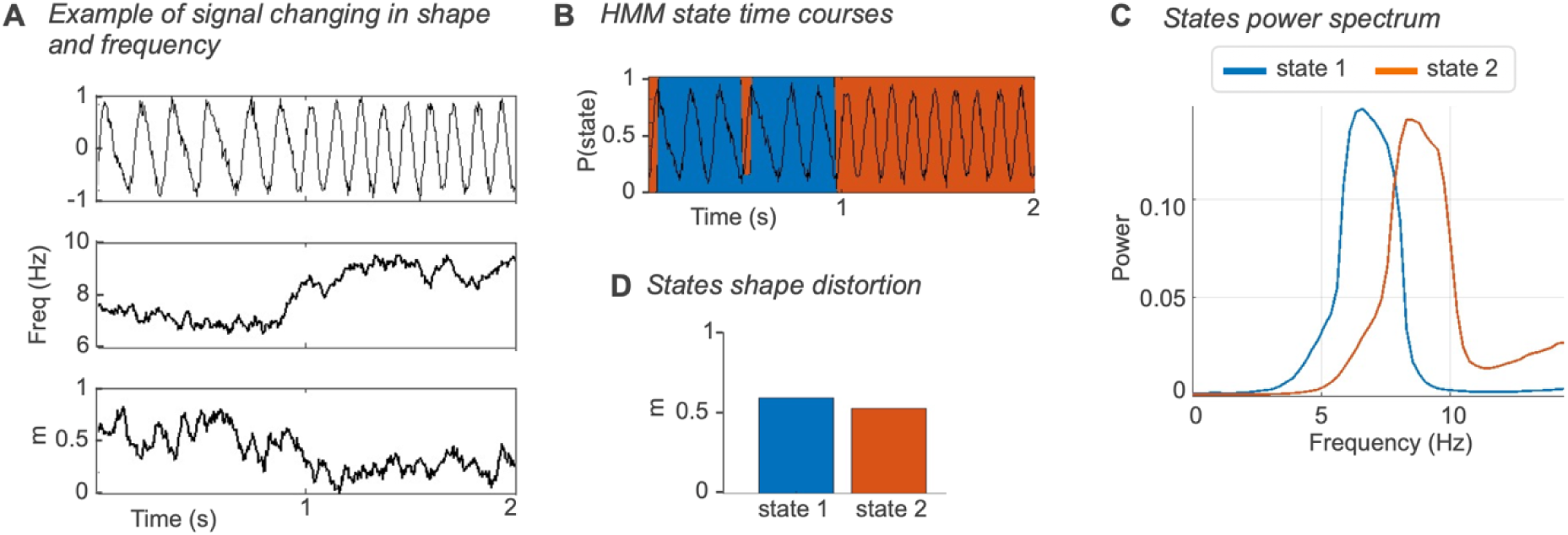
HMM analysis on synthetic data changing in shape and frequency independently. **A.** *Example of signal changing in shape and frequency*. Example of the synthetic signal (top panel), with its instantaneous frequency (middle panel) and the instantaneous shape distortion coefficient m (bottom panel). **B.** *Example of the HMM states time courses* on the signal plotted in **A. C.** *States power spectrum.* Power spectrum of the HMM states found on the signal in **A**. The states show different frequency peak. **D.** *States shape distortion.* The average distortion of the signal captured by each state. Each state captures a certain amount of distorted signal, meaning that the HMM state decomposition is driven mostly by frequency.

**Supplementary Figure 7:**
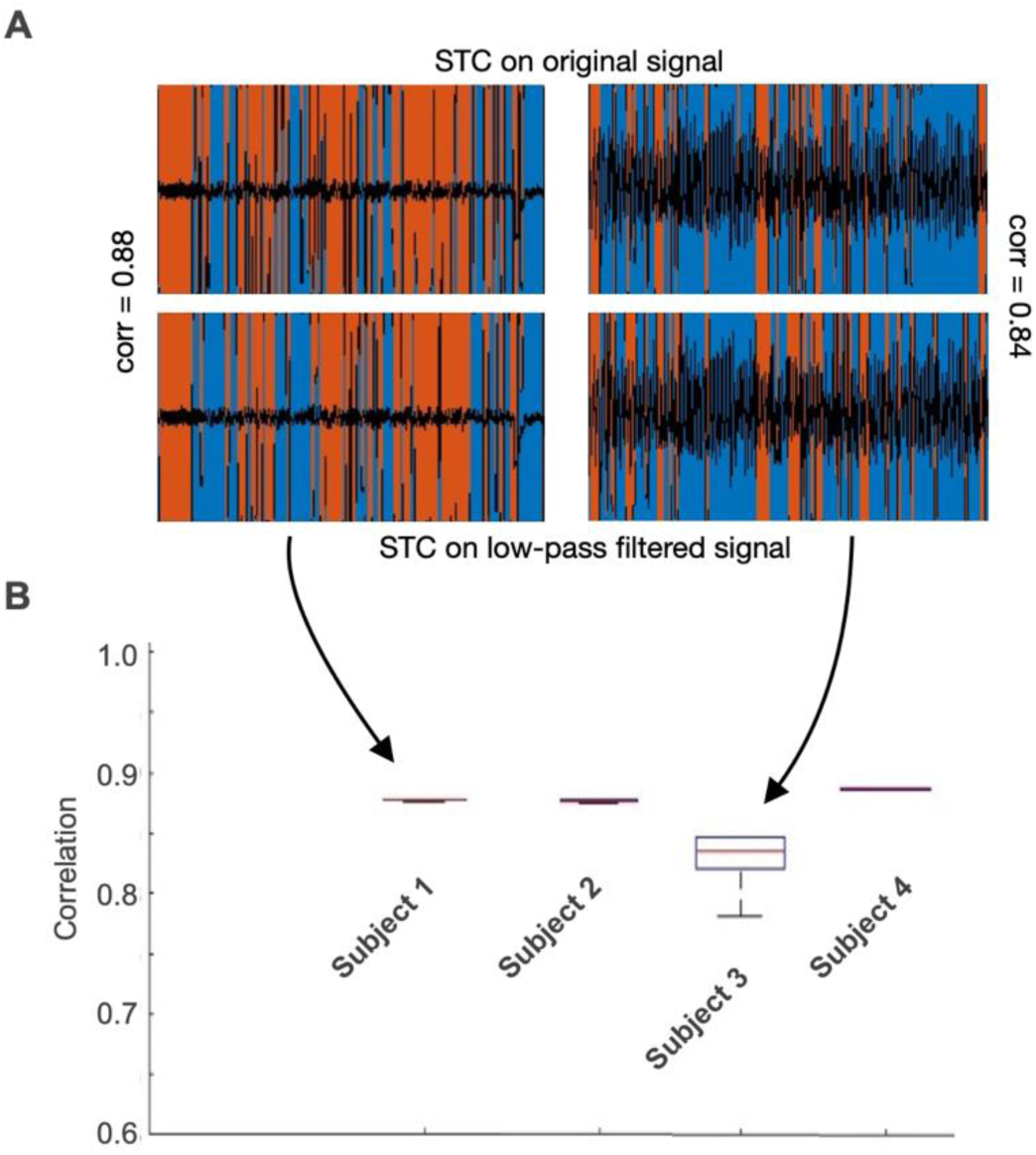
Correlation between the HMM run on the minimally pre-processed LFP signal and the HMM run on the low-pass filtered LFP signal. **A.** Two examples of HMM state time courses (STC) on the minimally pre-processed LFP data (top row) and on the low-pass filtered data (bottom row), and their correlation, for Subject 1 (highly correlated) and Subject 3 (relatively less highly correlated). **B.** Correlation between HMM runs on the minimally pre-processed signal and on the low-pass filtered signal, over 10 different runs. The HMM on the minimally pre-processed signal is run every time with random initial conditions. The HMM on the low-pass filtered signal is run by setting as initial conditions the HMM on the minimally pre-processed signal (to control for the inherent variability of stochastic inference of HMM). The red line in the box shows the mean, the box indicates the variance, and the eventual extra line indicates outliers.

**Supplementary Figure 8:**
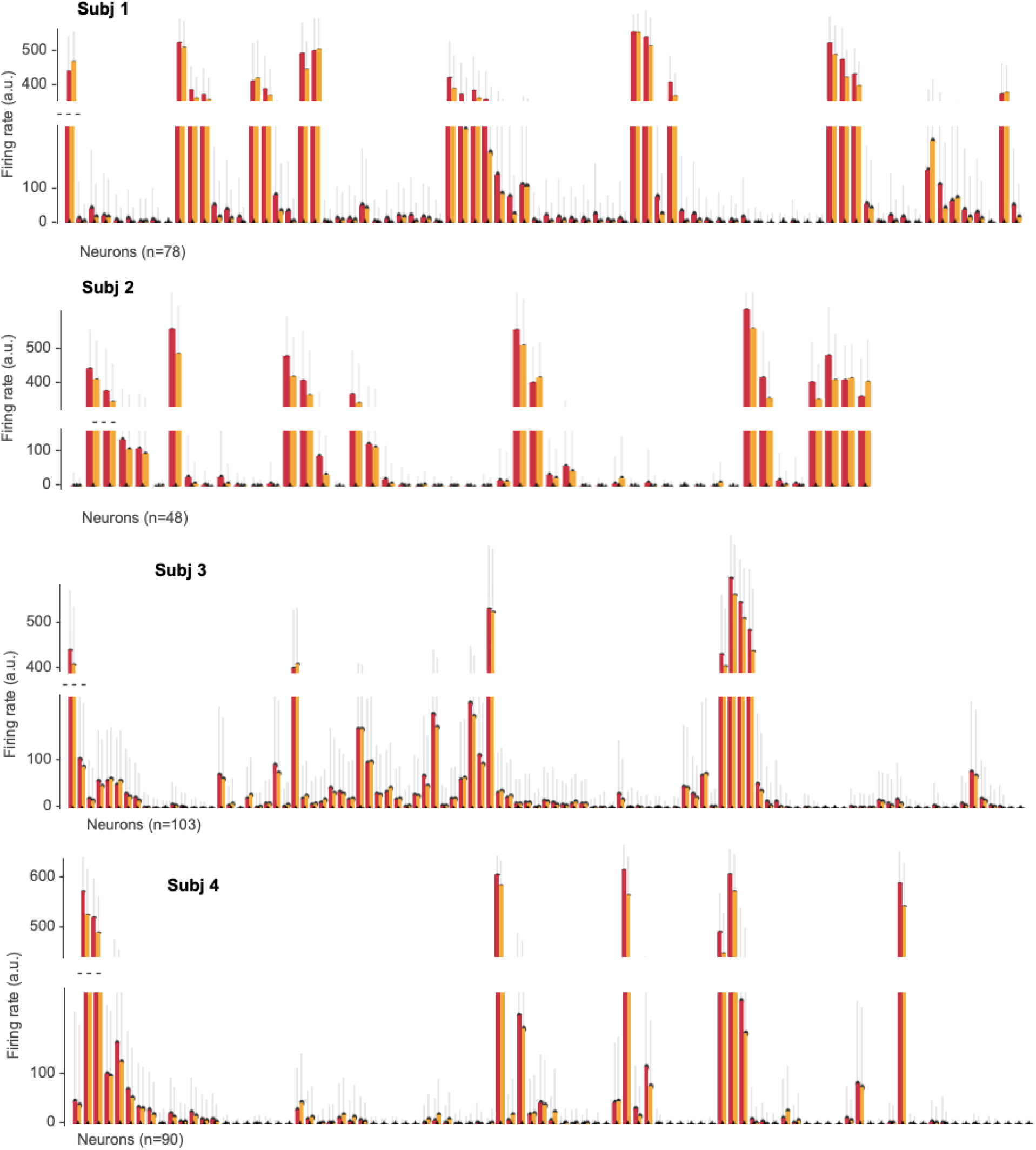
Neuronal average firing rate per state. Average firing rate and its standard deviation for each neuron during the different states. The HPHT (red) state involves higher neuronal firing rate, on almost every neuron.

**Supplementary Figure 9:**
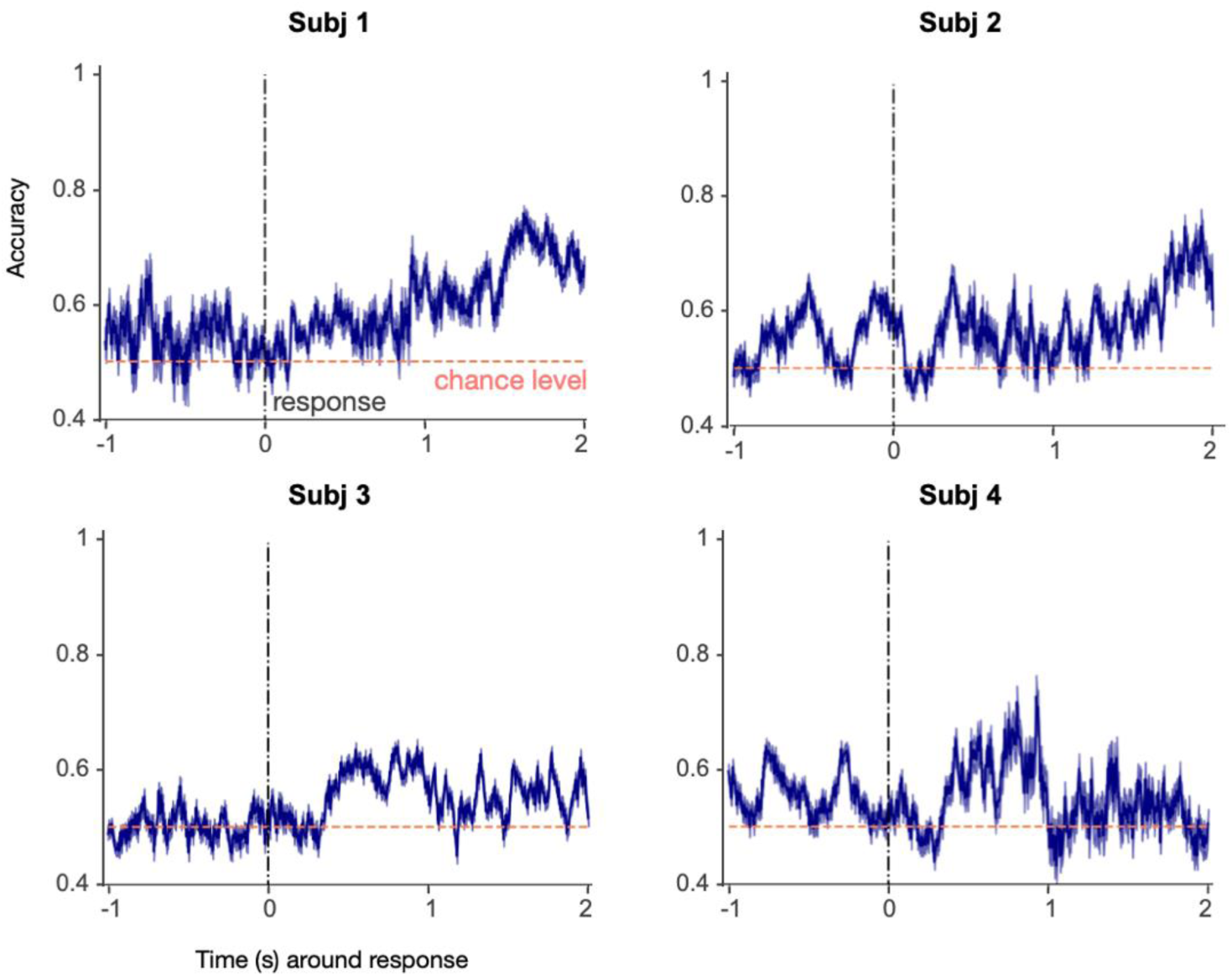
Prediction of state activation from the neural population activity. For each subject, we show the accuracy of decoding states activation from the neural population activity, standardised across neurons, per trial and per time point, to rule out that specific subpopulations of neurons are preferentially indexed by the states. The standardisation step is needed to be sure that the average firing rate, instead of the relative activation of the neurons, is not driving the predictions. This is because the HPHT state generally involves higher firing rates for most neurons. The prediction accuracy is averaged over 100 repetitions of a class-balanced decoding procedure, and is here reported within its 95% CI. The accuracy typically around chance level indicates that the states do not selectively recruit different neurons within the population.

**Supplementary Figure 10:**
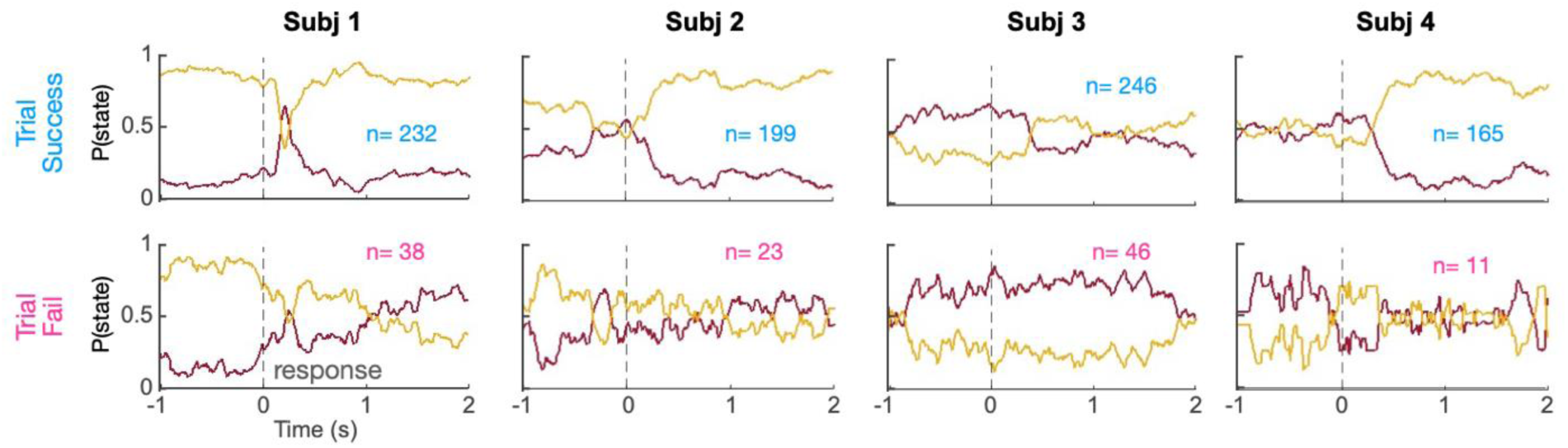
States distribution with respect to trial outcome. For each subject, we show the probability of each state being active around response, for trials that have been successful (top row) or failed (bottom row). The states activation probability is computed as an average across trials (n indicates the number of trials available), aligning trial data at response. Typically, the LPLT (yellow) state tends to be the most active during successful trials, while there is not a clear state dominance during the failed trials.

**Supplementary Figure 11:**
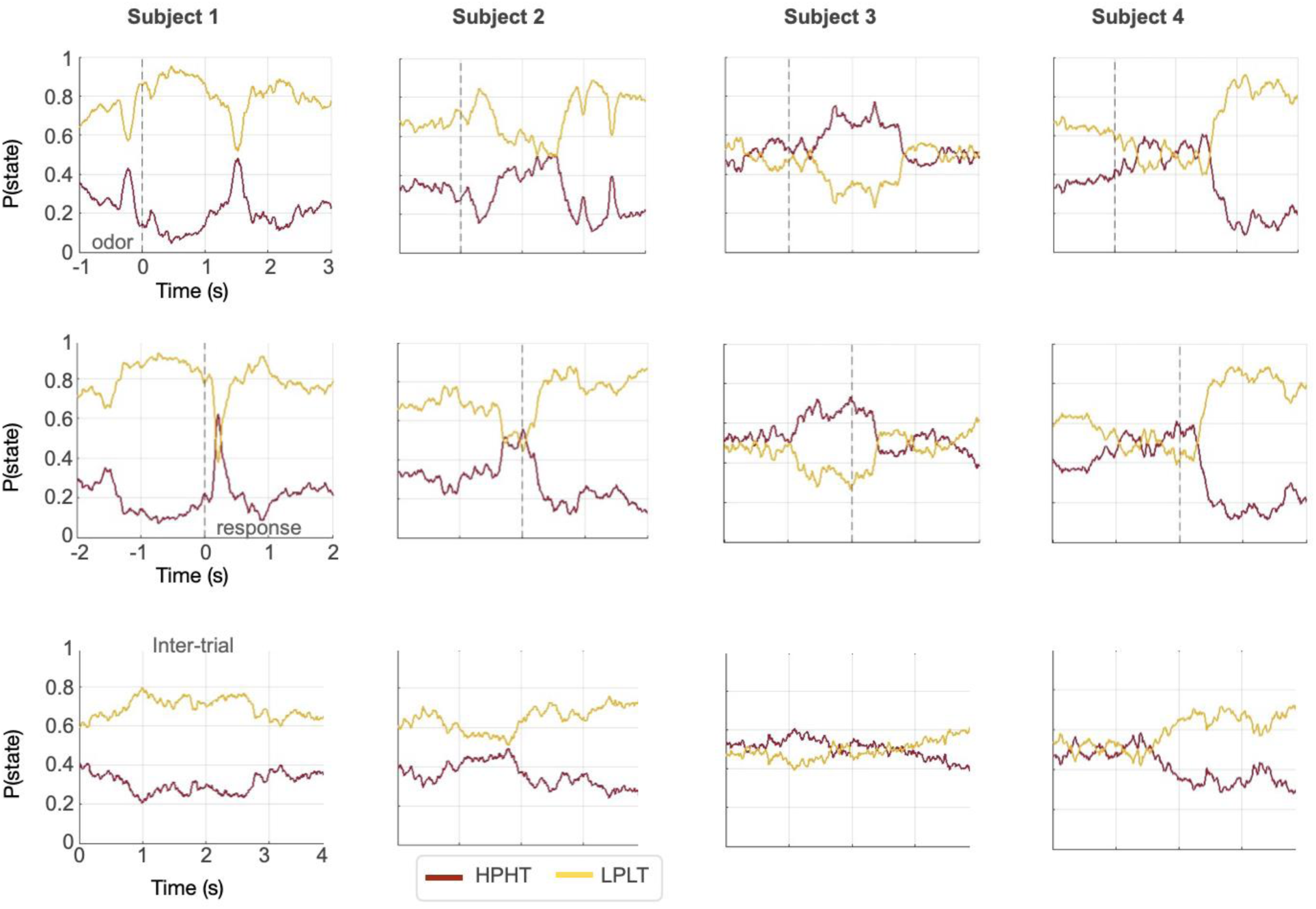
States distributions around task-relevant moments. Distributions around odor presentation (top row), response (middle row) and inter trial time (bottom row), for each subject.

**Supplementary Figure 12:**
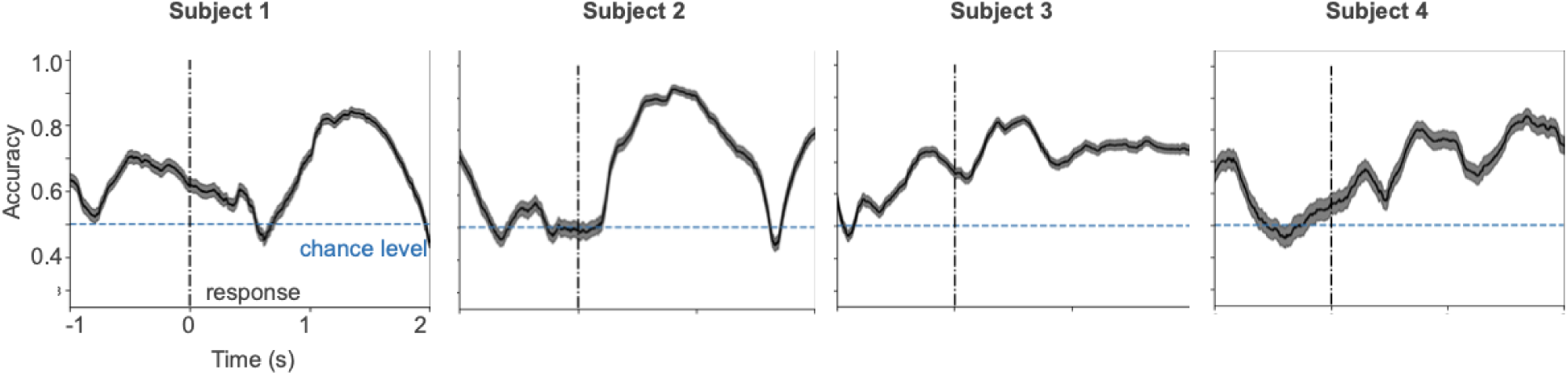
Predicting trial outcome from head position. For each subject, shown the prediction accuracy of trial outcome from Xp and Yp coordinates, within its CI of 95% as obtained with 200 bootstraps to ensure class balance. Decoding accuracy is high especially after response, likely reflecting the licking-induced movement of rewarded trials.

**Supplementary Figure 13:**
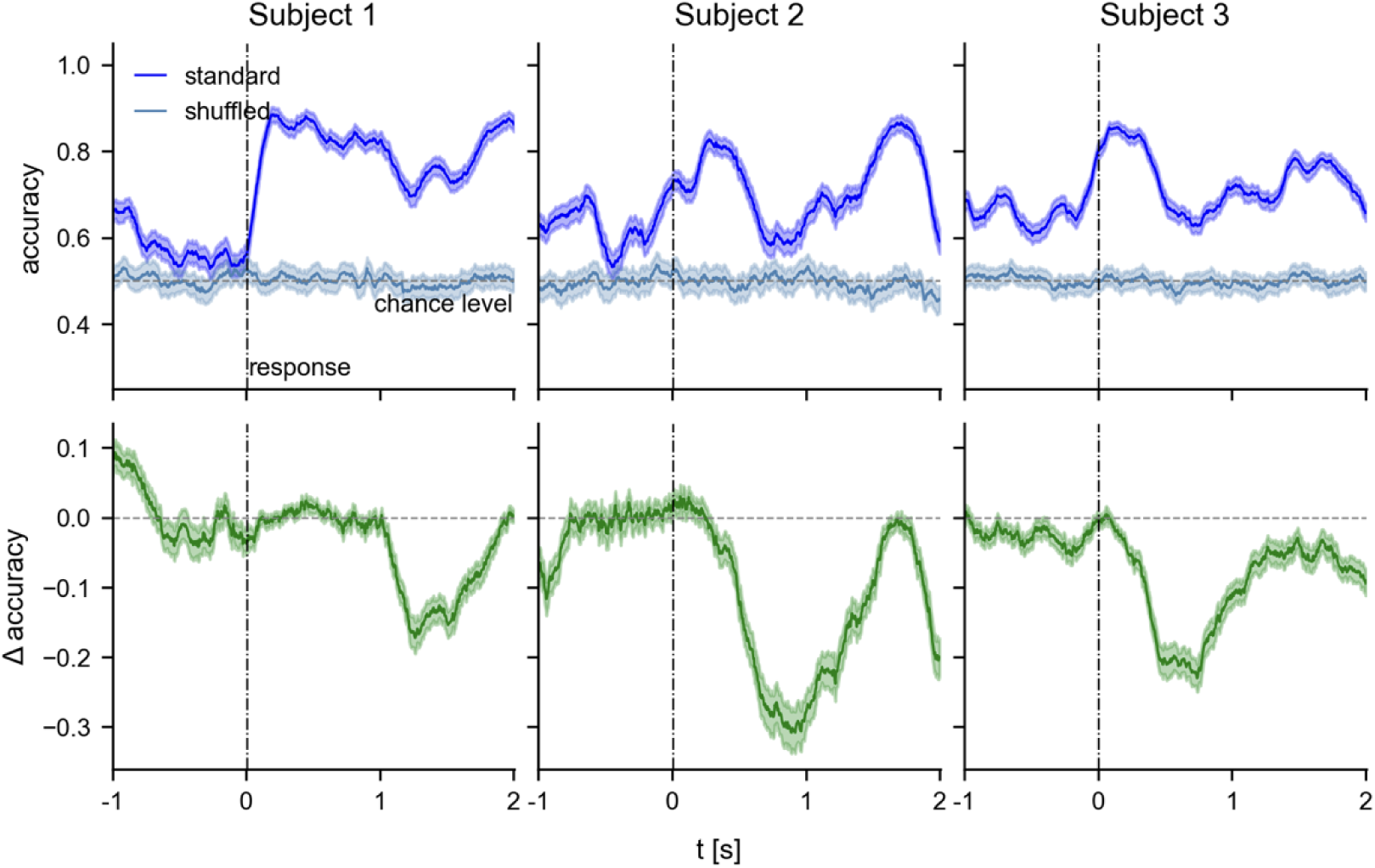
Standard and shuffled decoding from deconfounded neural activity. **Top row**: For each subject, shown the prediction accuracy of trial outcome from the neural activity deconfounded for head motion (Xp and Yp coordinates), within its CI of 95% as obtained with 200 bootstraps to ensure class balance. For comparison, the decoding is performed also on deconfounded neural activity with shuffled trial outcome labels (cyan, 200 permutations, 95% CI), as expected at chance level. **Bottom row:** the difference between standard decoding performed on the original neural activity and on neural activity deconfounded for head motion.

**Supplementary Figure 14:**
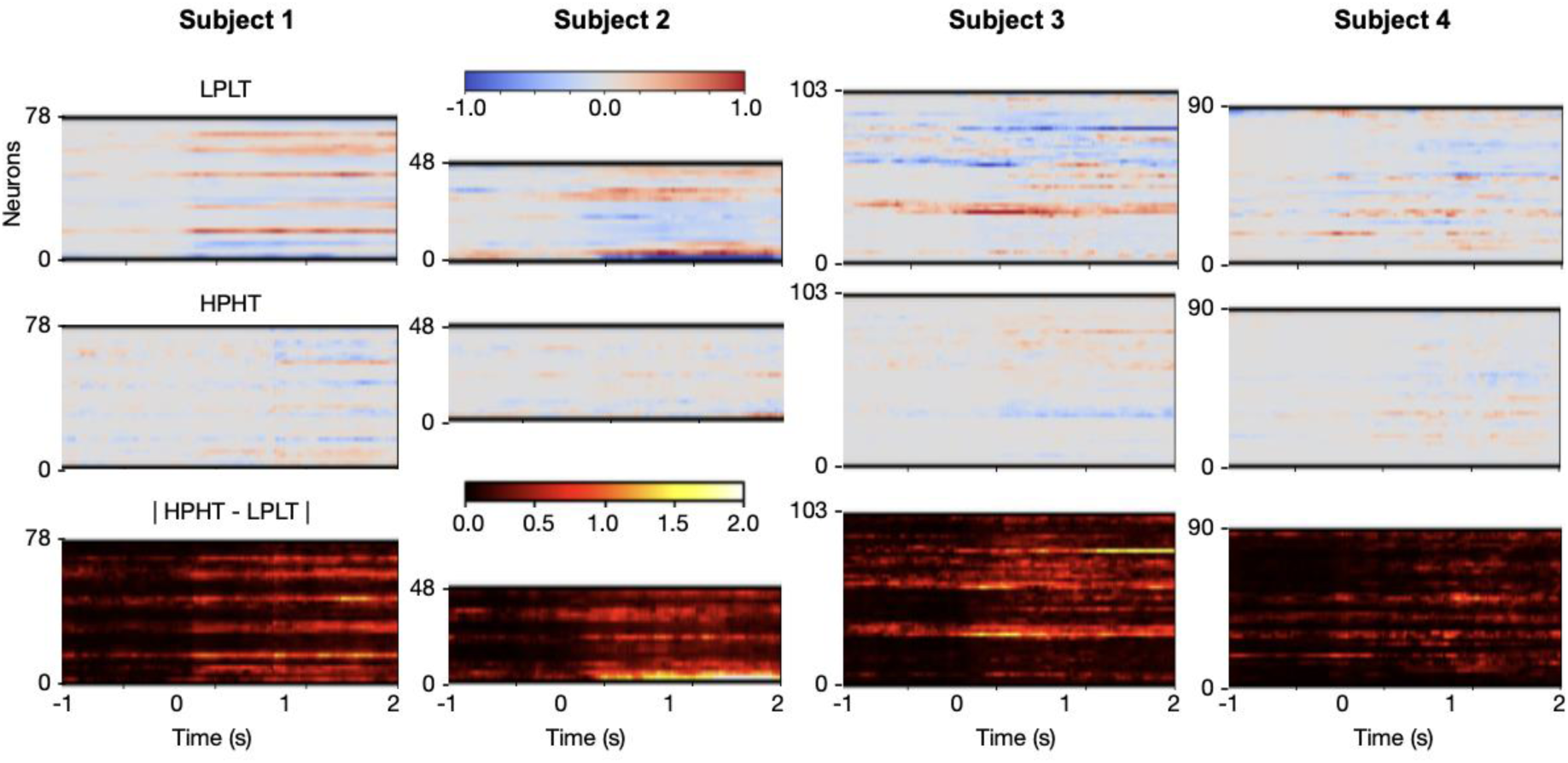
Neural activity representing success vs fail across states. The plots show the difference in neural activity averaged across trials for successful minus failed trials, for those trials in state LPLT (top row), and in HPHT (middle row). The bottom row shows the absolute difference of the activity in the two higher panels

**Supplementary Figure 15:**
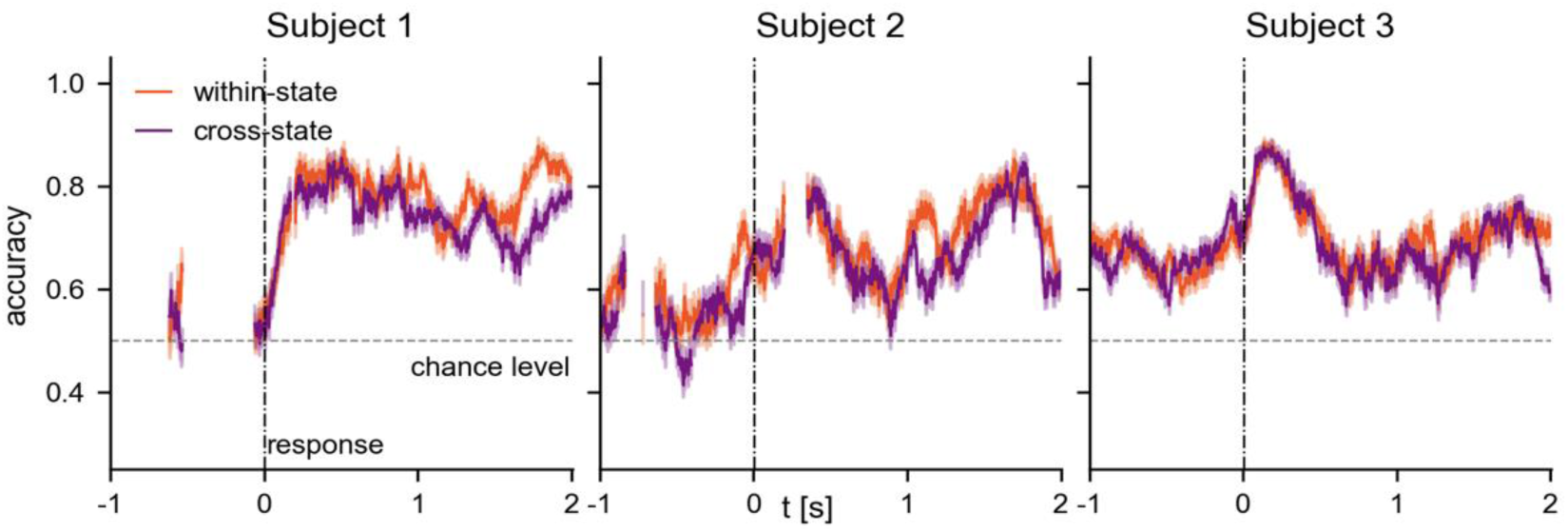
Within-state and cross-state decoding accuracy (signal filtered <100Hz). For each subject, shown the average decoding accuracy within-state (orange) and cross-state (purple) within their 95% CI (200 bootstraps to ensure class-balance), with the states obtained by running the HMM on the low-pass filtered signal (sharp filter at 100Hz). If the number of trials for each state at any point is lower than 5 per class, the decoding is not performed (gaps in the plot).

**Supplementary Figure 16:**
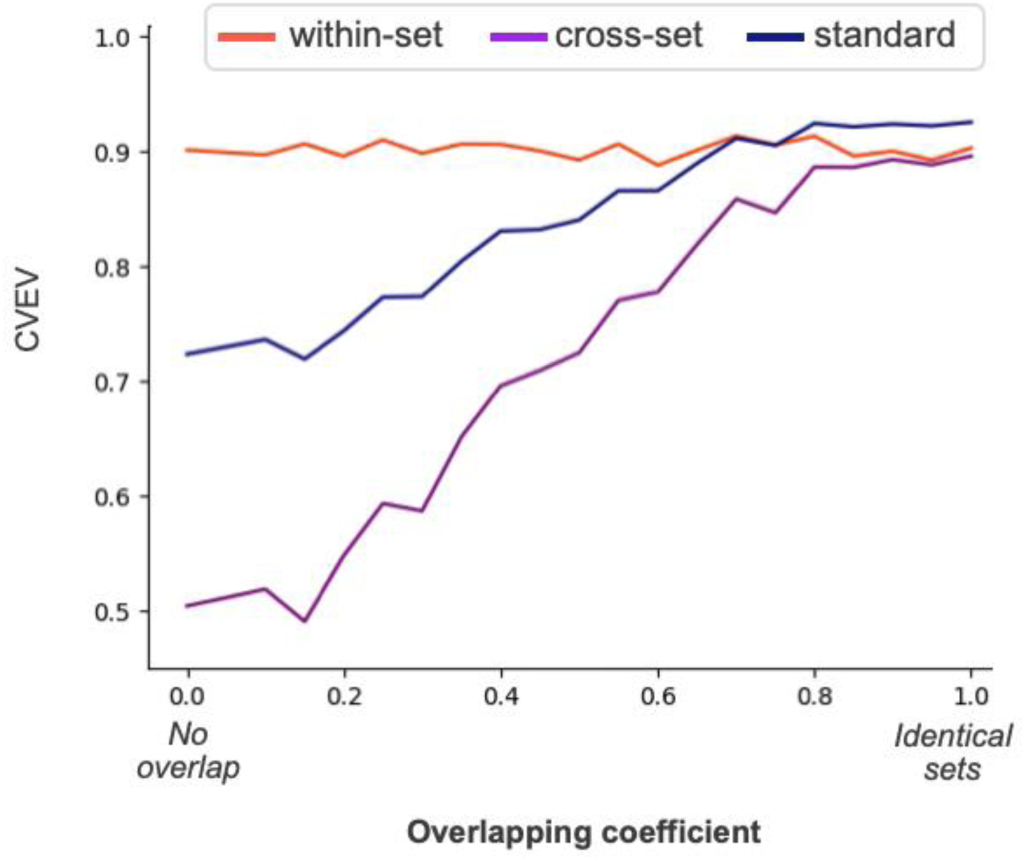
Decoding accuracy from simulated sets. The plot shows cross-validated explained variance (CVEV) of simulated sets, encoding information with varying degrees of overlap. Reported the CVEV of within-set, cross-set and standard decoding (i.e., decoding from both sets jointly, without distinguishing between the two), averaged over 500 simulated sets for each value of the overlap degree (20 values between 0 and 1). For values between 0 and 0.5 of the overlap degree (no overlap to 50% shared information encoded), within-set decoding works better than standard decoding and cross-set decoding performs much worse than both. For values between 0.6 and 1 of the overlap degree (1 indicating identical encoding weights for the two sets), standard decoding surpasses within-set decoding, and the cross-set decoding performance is increasingly higher. Between 0.7 and 0.8 of overlap degree lies the matching between these simulations and our experimental results.

**Supplementary Figure 17:**
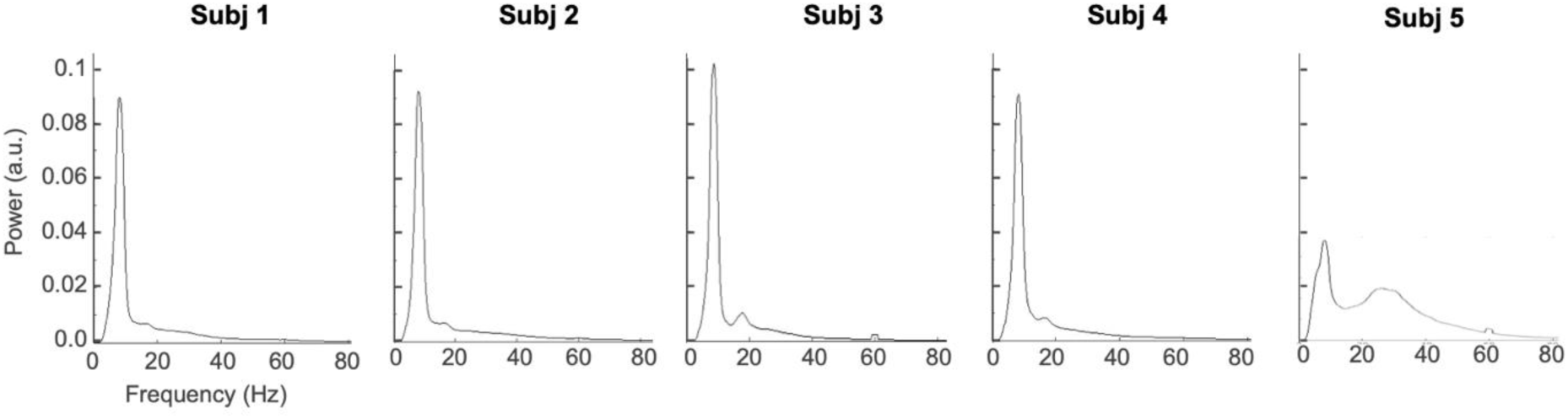
LFP data power spectrum of the 5 rats. Power spectrum of the 1^st^ principal component (PC) of the LFP data (high-pass filtered at 4Hz) for the five rats. Because of how different its power spectrum from the other rats was, Subject 5 was not included in the main report. We have performed all the analyses on this animal, reported in **Supplementary Figure 18**.

**Supplementary Figure 18:**
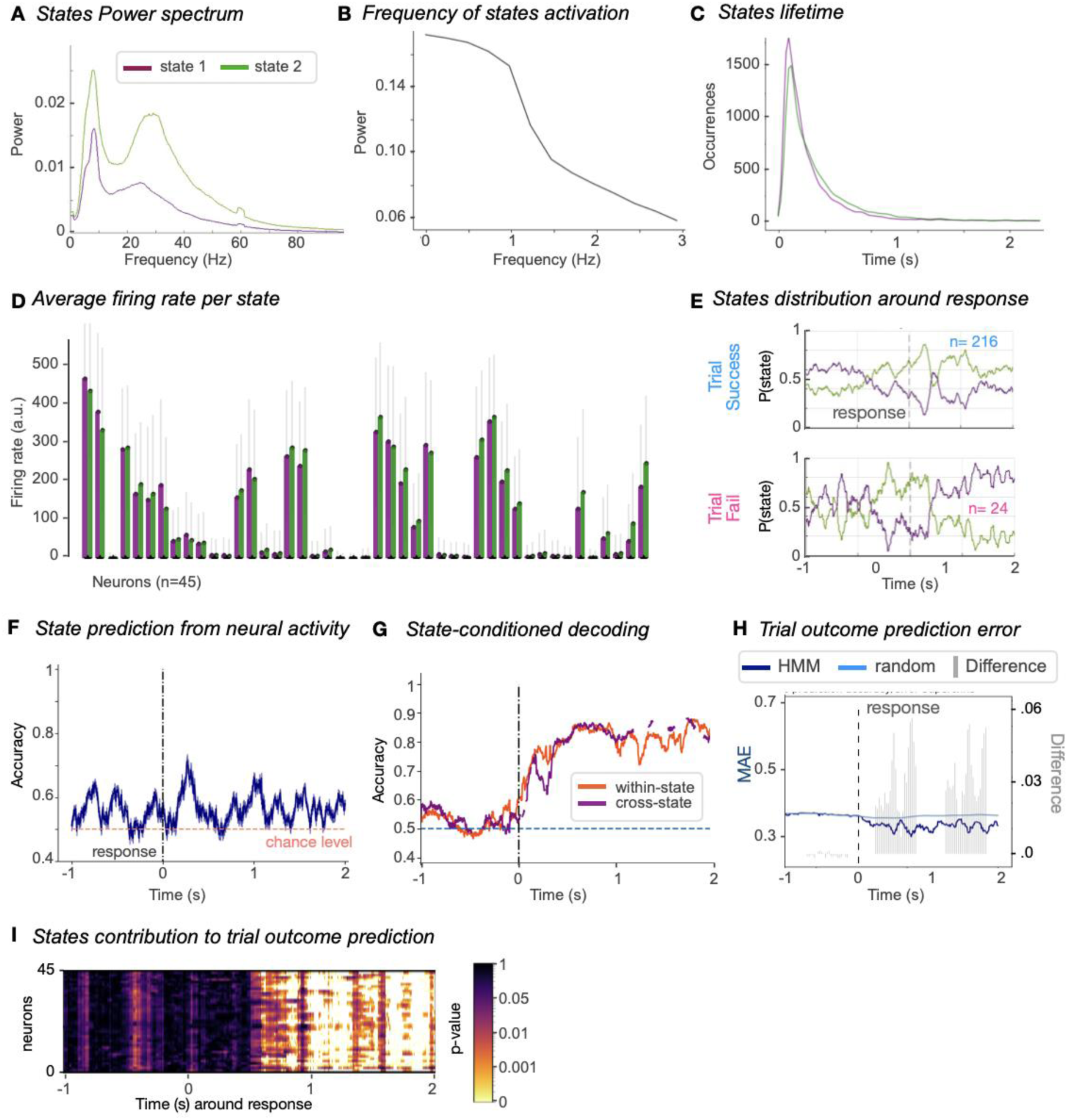
Analyses performed on subj 5. The HMM analyses were performed with the exact same settings as in all the other subjects. **A.** *States power spectra*. The states capture general differences in power across the frequency spectrum, but there is no specific LPLT and HPHT states. **B.** *States periodicity*. Similarly to all other subjects, the periodicity of the states peaks below 2 Hz, meaning at most two states switch per second. **C.** *States lifetimes*. Both states have a similar lifetime distribution. **D**. *Average firing rate per state*. Similarly to the other subjects, the state associated with higher power typically involves a higher firing rate across neurons. **E.** *State distribution around response*. The state involving higher power across the frequency spectrum is the most active during successful trials, while there is no clear dominance for failed trials. **F.** *Prediction of state activation from standardised neural activity*. Similarly to the other subjects, the prediction accuracy is typically around chance level, indicating that the state-neural activation relationship goes beyond the mere average firing rate. **G.** *State-conditioned decoding*. Within-state vs cross-state decoding accuracy. In some time points after response within-state accuracy is higher than cross-state. The points where cross-state decoding accuracy is missing indicate timepoints where the number of trials for one state is not enough to perform decoding. **H.** *Trial outcome prediction error* (state-neuron interaction analysis). Like in the other subjects, also here the HMM states help better predicting trial outcome after response. **I.** *States contribution to prediction of trial outcome*. Shows the significance of the improved prediction accuracy when adding the HMM states as regressors in the prediction of trial outcome.

## Notes

### Competing Interest Statement

The authors have declared no competing interest.

### Summary of Updates

Complete re-analysis of the data deconfounding for motion, new figures visualizing the effect described, new figures and analyses characterizing the HMM states

